# Rational positioning of 3D printed micro-bricks to realize high-fidelity, multi-functional soft-hard interfaces

**DOI:** 10.1101/2023.01.21.525002

**Authors:** M. C. Saldívar, S. Salehi, R. P. E. Veeger, M. Fenu, A. Cantamessa, M. Klimopoulou, G. Talò, M. Moretti, S. Lopa, D. Ruffoni, G.J.V.M. van Osch, L. E. Fratila-Apachitei, E. L. Doubrovski, M. J. Mirzaali, A. A. Zadpoor

## Abstract

Living organisms have developed design principles, such as functional gradients (FGs), to interface hard materials with soft ones (*e.g*., bone and tendon). Mimicking such design principles can address the challenges faced when developing engineered constructs with soft-hard interfaces. To date, implementing these FG design principles has been primarily performed by varying the ratio of the hard phase to that of the soft phase. Such design approaches, however, lead to inaccurate mechanical properties within the transition zone. That is due to the highly nonlinear relationship between the material distribution at the microscale and the macroscale mechanical properties. Here, we 3D print micro-bricks from either a soft or a hard phase and study the nonlinear relationship between their arrangements within the transition zone and the resulting macroscale properties. We carry out experiments at the micro- and macroscales as well as finite element simulations at both scales. Based on the obtained results, we develop a co-continuous power-law model relating the arrangement of the micro-bricks to the local mechanical properties of the micro-brick composites. We then use this model to rationally design FGs at the individual micro-brick level and create two types of biomimetic soft-hard constructs, including a specimen modeling bone-ligament junctions in the knee and another modeling the nucleus pulposus-annulus fibrosus interface in intervertebral discs. We show that the implemented FGs drastically enhance the stiffness, strength, and toughness of both types of specimens as compared to non-graded designs. Furthermore, we hypothesize that our soft-hard FGs regulate the behavior of murine preosteoblasts and primary human bone marrow-derived mesenchymal stromal cells (hBMSCc). We culture those cells to confirm the effects of soft-hard interfaces on cell morphology as well as on regulating the expression of focal adhesion kinase, subcellular localization, and YAP nuclear translocation of hBMSCs. Taken together, our results pave the way for the rational design of soft-hard interfaces at the micro-brick level and (biomedical) applications of such designs.

## 1. Introduction

Natural materials have developed smart design principles over millennia of evolution to interface materials with highly dissimilar mechanical properties (*e.g*., a hard material like bone and a soft material like cartilage or tendon) [1,2]. These structural interfaces, commonly known as functional gradients (FGs), exhibit specific mechanical property transition functions (*e.g*., linear, power, exponential) [3,4] and are present in a vast array of biological systems, including the squid beak [5], dentinoenamel junction [6,7], bone-soft tissue insertion [8–10], and byssal thread [11]. The development of advanced materials with enhanced, mutually exclusive mechanical properties (*e.g*., strength and toughness) are often inspired by such biomimetic design principles [12] to address challenges associated with the arising stress concentrations and the mismatch between the load-carrying capacities of both materials [13–16].

Recent progress in polymer-based multi-material additive manufacturing (AM, known as 3D printing) [17,18] has enabled the realization of FGs through several processes, such as material extrusion [19–23] and material jetting [24–26]. Among those techniques, voxel-based material jetting provides unparalleled freedom to design complex structures, thanks to its hallmark drop-on-demand capability [27,28]. Voxel-based design of soft-hard interfaces is then akin to the positioning of soft and hard micro-bricks with side lengths of, say, 40 μm next to each other to create a specific transition zone between 100% hard and 100% soft micro-bricks. Different variations of this technique have already been used to generate hierarchical and graded constructs with improved strength and toughness [29–31]. At the macroscale, however, we usually do not care about the exact organization of micro-bricks but would, instead, like to realize transition zones with specific variations in the mechanical properties. The studies performed to date have mainly analyzed such transition zones in terms of the ratio of the number of hard micro-bricks to that of soft micro-bricks without considering all the possible permutations of hard and soft micro-bricks [32–34]. Moreover, the relationship between the numbers of hard and soft micro-bricks and the realized macroscale properties is often assumed to be linear, neglecting its highly nonlinear nature. It has already been shown that such assumptions can lead to inaccurate estimations of effective mechanical behavior [35–37].

Here, we aim to establish nonlinear models that relate the positioning of micro-bricks to the actual values of the elastic modulus within the transition zone of FG soft-hard interfaces. We then use these models to create FG soft-hard interfaces with multiple types of functionalities. Our methodology combines experimental tools to characterize FGs through nanoindentation experiments at the microscale, quasi-static tensile tests analyzed with digital image correlation (DIC) at the macroscale, with detailed finite element models at both scales. We showcase the applications of such FG soft-hard interfaces by: 1. rationally designing the soft-hard interfaces of two types of 3D printed biomimetic constructs, and 2. demonstrating that such transition zone can be used to regulate cell behavior. The biomimetic constructs include a bone-ligament junction of the knee and the nucleus pulposus-annulus fibrosus interface of an intervertebral disk. As for the second application, property-based FGs are shown to direct the migration (*i.e*., durotaxis) and potential differentiation of living cells [38–42]. Here, we demonstrate that our micro-brick positioning approach can be used to regulate the behavior of murine preosteoblasts and human bone marrow-derived mesenchymal stromal cells (hBMSCs). Toward this aim, we analyzed the morphological differences, FAK expression, subcellular localization, and YAP nuclear translocation of primary hBMSCs across graded and non-graded specimens.

## 2. Results and discussion

### 2.1. Material characterization and modeling

Using voxel-based AM technology, we 3D printed two types of prismatic-shaped specimens with a linear gradient of hard material volume fraction (*ρ*) projected into their volume (Figure 1A-C). We used VeroCyan^™^ (Stratasys^®^ Ltd., USA) UV-curable photopolymer as the hard phase for both specimens. For the soft material, however, we assigned Agilus30^™^ Clear (Stratasys^®^ Ltd., USA) to the first specimen type and MED625FLX™ (Stratasys^®^ Ltd., USA) to the second specimen type. We tested these specimens using a nanoindentation (NI) protocol [32,43] which allowed us to interrogate multiple locations within the FG transition zones, revealing the entire elastic behavior achievable by these composites (Figure 1B-E). Furthermore, the mechanical properties of several representative volumetric elements (RVEs) extracted from these specimens were simulated using the finite element method (FEM), which indicated a good agreement between the simulations and experimental results (Figure 1D-E).

**Figure 1.**
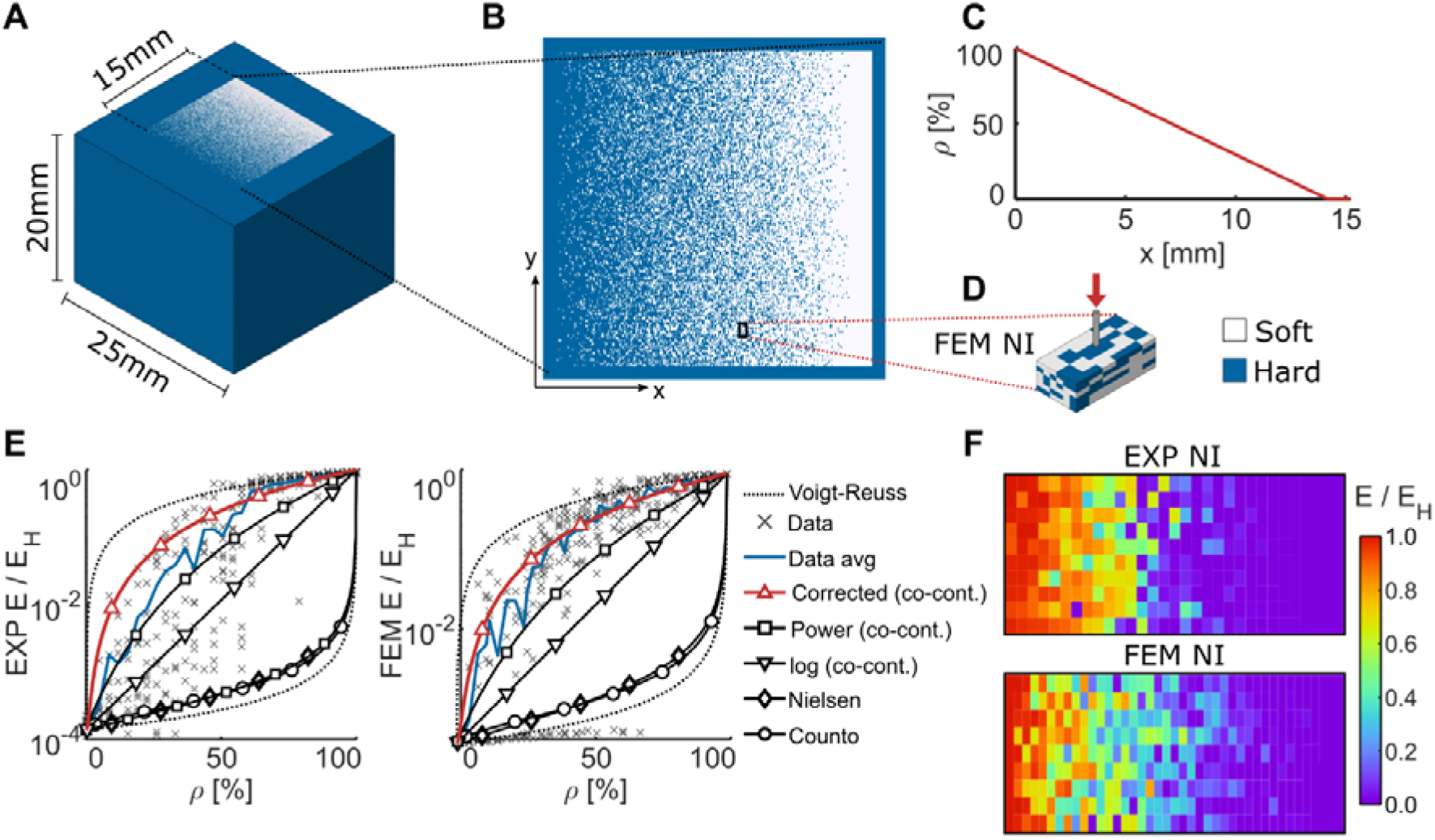
A) The design of the 3D printed specimen used for testing the properties of voxel-based composites via nanoindentation. B) A representative binary image of the nanoindentation specimens with a gradient of material properties projected onto their geometry. C) The *ρ* function applied to the specimen for measuring the properties of these composites across their entire property space. D) A representative volumetric element of a subsection of the nanoindentation specimens used to create FEM models of nanoindentation. E) The normalized elastic modulus *vs. ρ* and the corresponding average response (in blue) measured (EXP) through nanoindentation and predicted by computational models (FEM). These values and their associated trend lines are compared with several existing models for these composites. The modified co-continuous model was found to be the most accurate. F) The heatmap plots of the elastic moduli measured with nanoindentation and the corresponding FEM predictions across the FG specimens with a linear variation of *ρ*.

On average, the elastic modulus of the hard material, *E_H_*, was 1994.7 (±74.89) MPa. The elastic moduli of the soft materials, *E_S_*, were 0.507 (± 0.171) MPa and 5.4282 (± 2.72) MPa for Agilus and MED625FLX, respectively. For the linear gradient of *ρ*, both the experiments and simulations showed high variations in the local elastic response (Figure 1E and Figure S1A-B of the supplementary document). The heterogeneous nature of the composites at the micro-brick (*i.e*., voxel) scale caused these variations, which is the length scale probed during the NI tests (Figure 1F). Regardless of the local variations, the average response of both types of specimens showed a nonlinear transition of elastic modulus across the gradient, which contrasted with the linear transition in *ρ*. Moreover, the observed behaviors of both types of specimens and their corresponding FEM estimations were remarkably similar (Figure S1C of the supplementary document). These two observations confirmed the validity of the obtained data and allowed us to use them as input for the mechanical characterization of such 3D printed micro-brick arrangements.

Most of the predictions made by the classic models of particle-reinforced composites did not match the elastic response observed during our experiments and simulations. These models included those proposed by Nielsen [44], Counto [45], and the simplified power and logarithmic co-continuous models proposed by Davies [46] (Section S1 of the supplementary document). Among these models, the simplified power-based co-continuous model was the most accurate. This observation is in line with the findings of a recent study performed on non-voxelated specimens with homogeneous distributions of *ρ* [43]. The residual plots of this model, however, highly increased for most *ρ* values (Figure S1 D-F of the supplementary document). These high errors indicate that this version of the co-continuous model is insufficient for capturing the elastic behavior of the voxel-based composites studied here. We, therefore, generalized the power law-based co-continuous model as:

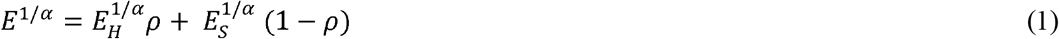

where *E* is the elastic modulus of the composite material. Similarly, *α* is a power law coefficient that determines the nonlinear behavior of the composite and depends on the geometrical arrangement of the micro-bricks, the ratio of the elastic modulus of both phases (*E_H_*/*E_S_*), and the particle joint probability function of the micro-brick arrangements [39]. After fitting this parameter with a bi-square nonlinear regression algorithm, we obtained *α* = 1.95 (95% confidence interval, C.I. = 1.82 - 2.1) and *α* = 1.86 (95% C.I. = 1.75 - 1.97) for the experiments on AgilusClear and MED625FLX, respectively. Similarly, we obtained *α* = 1.93 (95% C.I. = 1.83 - 2.04) for the FEM simulations. The residual of different models strongly depended on *ρ*. For *ρ* < 25%, higher residuals were observed, highlighting the complexity of capturing the behavior of composites when their mechanical behavior is dominated by the soft phase. Nevertheless, the nonlinear model proposed here achieved lower residual values across the entire design space of the micro-brick arrangements (Figure S1 D-E of the supplementary document) as compared to the simplified power and logarithmic co-continuous models. We, therefore, proceeded to the evaluation of the modified co-continuous model by designing FGs using the direct design of local elastic properties instead of designing the ratio of hard micro-bricks to that of soft micro-bricks.

### 2.2. Property-by-design of FGs

Generating three FGs with three different transition functions enabled us to evaluate the precision of the model (Equation 1). These FGs had linear (*E_lin_*(*x*)), step-wise (*E_ste_*(*x*)), and sigmoidal (*E_sig_*(*x*)) functions (Figure 2), which were 3D printed using VeroCyan and AgilusClear. To generate their equivalent *ρ*(*x*) functions, we used the inverse of Equation (1), which has the following form:

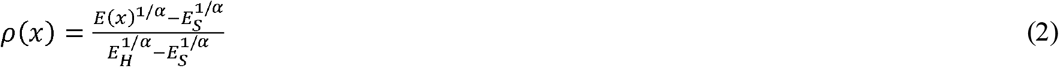

where *E*(*x*) is the desired FG function. For simplicity, we chose *α* = 2.0 to design these FGs because this value was within the 95% C.I. of the nonlinear fitting results obtained for both experiments and simulations.

**Figure 2.**
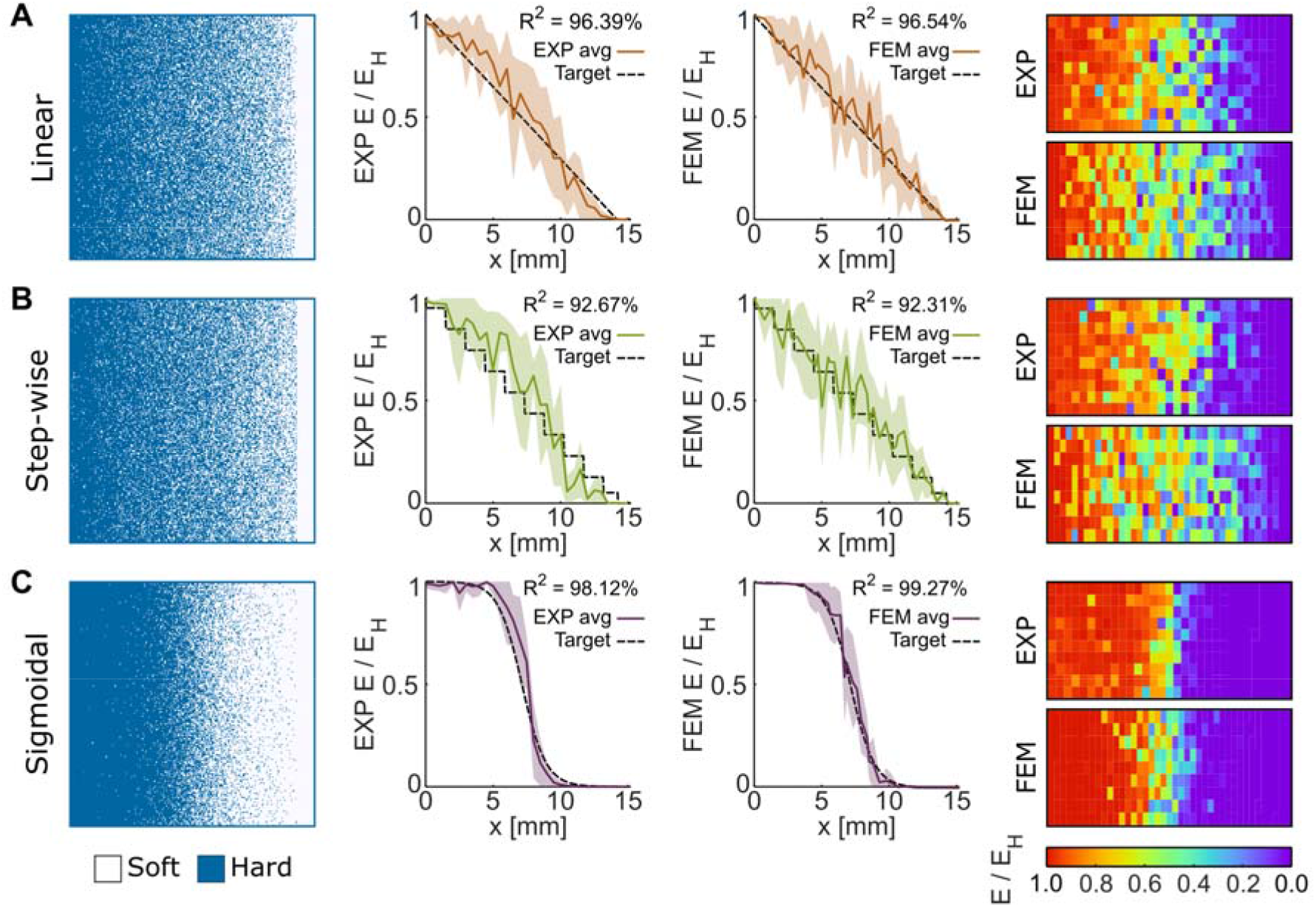
The measured and FEM-predicted nanoindentation results for various FG designs according to the modified co-continuous model. These tests were performed for the FGs with linear (A), step-wise (B), and sigmoidal (C) variations in the elastic modulus.

Similar to FGs with a linear *ρ*, we observed high variations in both experimentally obtained and FEM-predicted values of the local elastic moduli (Figure 2 A-C). The average behavior of the FGs was, however, highly correlated with the target elastic modulus functions (*i.e*., *R*^2^ > 92%), particularly for the sigmoid gradients (*i.e*., *R*^2^ > 98%, Figure 2C). Despite the higher number of estimation points for the simulations (*i.e*., 1405 simulation points per FG and 320 experimental points per FG), the average response and standard deviation of both types of characterization techniques were similar. Although many factors can complicate the NI testing of polymeric materials (*e.g*., adhesion and viscoelasticity), the strong similarity between NI and FEM suggests that the heterogeneous nature of the micro-brick composites is the main cause for the high standard deviations observed here. The mean trendlines of the elastic modulus of all the measurement groups resembled their corresponding designs except for the step-wise FG group, which despite having a high coefficient of determination (*R*^2^ = 92.67%), substantially deviated from its design function. The deviations observed in the step-wise FG group can be explained by the fact that the size of each step in that group was smaller than the observed variations in the local elastic response.

The NI experiments and their corresponding FEM simulations probed the properties of individual voxels and their close neighbors at the micrometer length scales. To assess how the microscale measurements relate to those performed at larger scales, we performed quasi-static tensile tests that measured the mechanical response of the specimens across the entire FG. We used digital image correlation (DIC) during those tests to measure the full-field strain distributions. We modeled the FGs as systems of linear springs subjected to tensile loading.

To obtain the mesoscale spatial variation of the elastic modulus from the experimentally measured strains, we assumed that the local elastic modulus value along the *x*-coordinate, *E*(*x*), is equivalent to the slope between the normal stress applied to the system (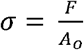, *A_o_* = 32.512 mm) and the average longitudinal (engineering) strains (*ϵ_p,avg_*(*x*)) of each cross-sectional layer of the FG (*E*(*x*) = *σ*/*ϵ_p,avg_*(*x*)).

The stresses measured for four groups of the tensile test specimens (*i.e*., power-law, linear, step-wise, and sigmoidal) monotonically increased with the applied strain (Figure 3B). The sigmoidal design exhibited the most compliant response. The local distributions of the elastic modulus were determined by the underlying design functions (*R*^2^ > 93%) (Figure 3C-F). Similarly, the effective elastic moduli, 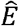, which is the equivalent modulus of the entire interface calculated from each *E*(*x*) value along the gradient, strongly correlated with the elastic moduli measured from the general stress-strain curves, *E_G_* (*R*^2^ = 95.76%, Table S1 of the supplementary document). Despite their considerable standard deviations, the average elastic modulus across the linear, power, and sigmoid gradients followed their target elastic modulus functions, validating the possibility of generating accurate FGs. The mean *E*(*x*) of the step-wise FG specimens, however, deviated from their corresponding design function. In fact, only the most compliant steps of these gradients were discernible. Two factors might have caused the absence of stiffer steps. First, the 6 times larger facet size of the DIC recordings as compared to the micro-brick size has likely led to the averaging of strains in the regions where sharp step transitions were present, blurring the measured step feature. Second, partial resin mixing at the interface between the micro-bricks may have resulted in a gradual transition of the elastic properties across the steps, similar to what other studies have suggested [37,43,47,48]. In contrast, none of these effects were present when simplifying the outcome from the FEM estimations into systems of linear springs. The obtained mechanical properties of the simulations showed the highest coefficients of determination in this study (*R*^2^ > 99 %), and the shapes of the gradients followed the expected gradient shapes. These results confirm that the deviations in the experimental measurements were due to the imaging resolution and potential material mixing effects. Therefore, these quasi-static tensile test experiments confirm that the presented model allows for the adjustment of the actual macroscale properties of voxel-based 3D printed FGs.

**Figure 3.**
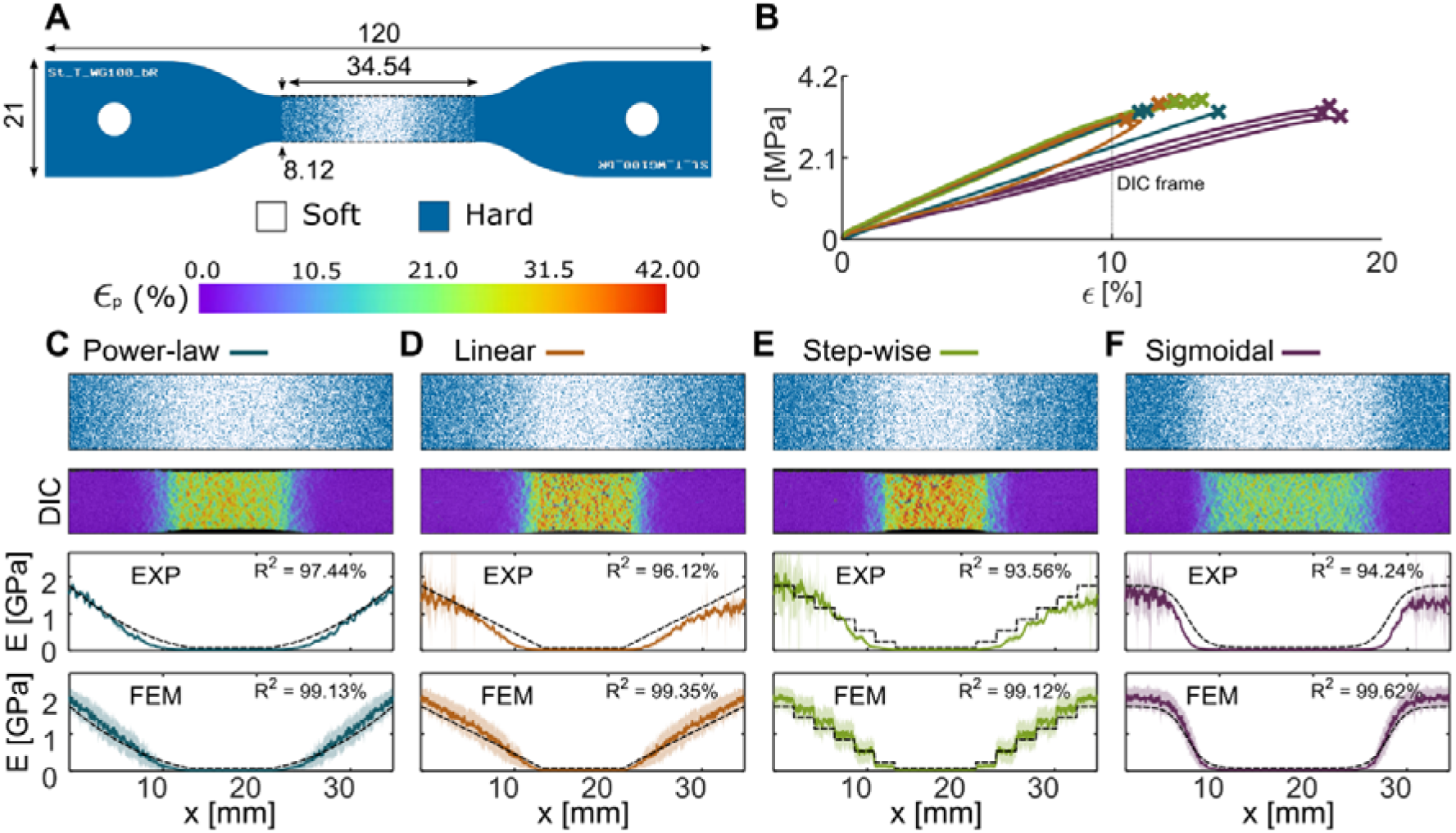
Additional FGs with different elastic modulus transition shapes, which were designed with the modified co-continuous model after idealizing the tensile test specimens under uniaxial deformations as systems of linear springs. A) A representative binary image of the tensile test specimens (out-of-plane thickness = 4 mm). The gauge region was designed with four symmetric FGs. We performed DIC measurements and FEM estimations to obtain the local deformations along the gradients. B) The stress-strain curves of the experiments. The FGs included those with power-law (C), linear (D), step-wise (E), and sigmoidal (F) changes in the elastic modulus.

### 2.3. Tough biomimetic structures

We used the modified co-continuous model proposed and corroborated in the previous sections to explore the applications of FGs in the design of clinically relevant biomimetic structures. First, we considered the challenging problem of interfacing soft and hard tissues, such as ligaments and bones, tendons and bones, and cartilage and bone. Toward this end, we designed two different systems of knee ligaments and performed quasi-static tension experiments and FEM simulations (Figure 4A). In the first group, we incorporated a sigmoidal FG into the design of each ligament-bone connection (Figure 4B and Figure S3B of the supplementary document). In the second group, however, we simply connected the soft and hard phases, effectively implementing a step function. The second group served as the control group (Figure 4C). The choice of sigmoidal functions was motivated by the results obtained in the above-presented tensile experiments and the fact that the strain distributions of these transition functions indicated a smooth transition between the hard and soft phases.

**Figure 4.**
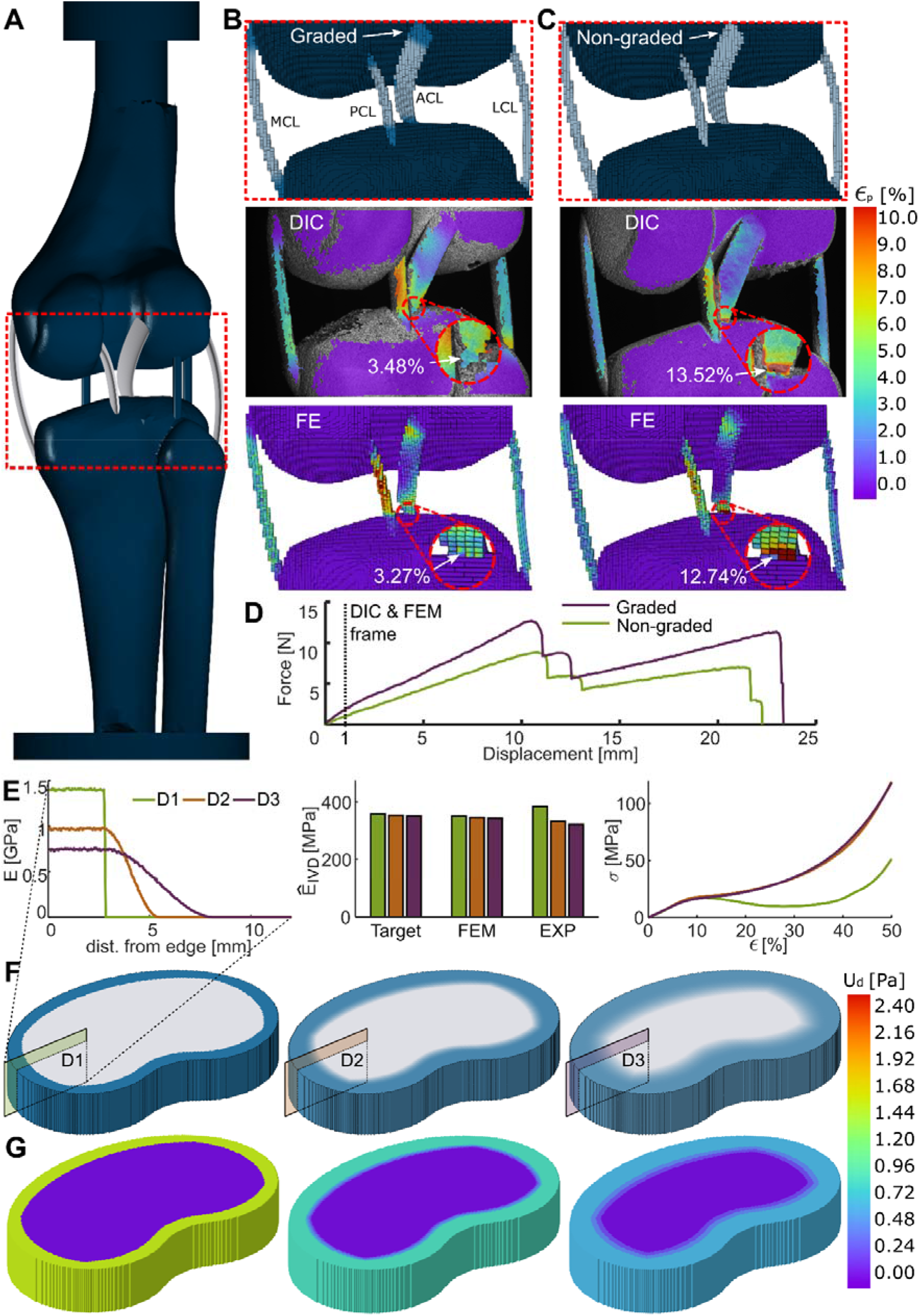
A case study where we implemented several strategies of functional grading to design biomimetic devices with enhanced strength and toughness. A) The design of a knee ligament system. B) graded and C) non-graded versions of this knee system were defined to study the failure mechanisms of the biomimetic bone-ligament connections through DIC measurements and FEM simulations. D) The force-displacement curves of both graded and non-graded designs. E) The different sinusoidal transition functions defined for each IVD design together with the resulting mechanical properties and stress-strain curves. F) The representative renders of each design after projecting the elastic modulus FGs onto each lamella of the IVD. G) The elastic strain energy density distributions resulting from the FEM simulations performed for all designs. The specimens were subjected to quasi-static compression.

The non-graded (*i.e*., control) design exhibited substantial strain concentrations at their soft-hard ligament interface, particularly within the ACL region (Figure 4C). This early onset of strain concentrations resulted in non-critical cracks for low displacements (Figure S3B of the supplementary document). Furthermore, the FEM predictions revealed that shear deformations at the ligament-bone interface cause the inadequate performance of the non-graded design (Figure S3C of the supplementary document). In contrast, the graded design showed lower strain values at the ligament-bone interfaces, indicating an improved distribution of stresses that led to a higher ultimate load before failure (Figure 4B). Moreover, the FEM simulations showed no substantial shear deformations in the graded system. These positive effects caused by the presence of FGs were reflected in the force-displacements curves of these experiments (Figure 4D). The graded structure was 1.3 times stiffer (*i.e*., *K_g_* = 1.09 N/mm *vs. K_ng_* = 0.84 N/mm), 1.44 times stronger (*i.e., F_max,g_* = 12.7 N *vs. F_max,ng_* = 8.82 N), and 1.55 times tougher (*i.e., U_g_* = 180.66 mJ *vs. U_ng_* = 116.77 mJ) than the non-graded design. It can, therefore, be concluded that decreased stress concentrations at soft-hard interfaces and reduced shear deformations improve the overall mechanical performance of the biomimetic FG design as compared to a non-graded design.

The second biomimetic, clinically relevant construct was an intervertebral disc (IVD) with rationally designed elastic properties (Figure 4E). Although similar bioinspired structures have been introduced in the literature [20], the gradient strategy applied in that study consisted of a step-wise FG to transfer the failure mode from the nucleus pulposus (NP) to the edge of the annulus fibrosus (AF). Their applied design methodology, however, resulted in a lower toughness as compared to that of non-graded designs. To overcome this issue, we assumed that the vertical deformations of an IVD under compression occur at the same rate across its surface and that the construct fails once the lamella with the lowest ultimate strain fracture. For soft-hard micro-brick composites arranged in parallel as NP and AF, this failure will typically occur in the region with the highest number of stiff micro-bricks (*i.e*., highest *ρ* value). We hypothesized that implementing an FG transition zone will reduce the interface stresses between the NP and AF. Moreover, we adjusted the maximum *ρ* value within the AF to be high enough to enable the construct to withstand physiological loads while remaining as low as possible to maximize its potential to store strain energy. Based on these strategies, we defined different transition functions within IVDs to increase their overall toughness while maintaining the same effective elastic response.

To demonstrate the design freedom provided by the micro-bricks, we designed three types of specimens with three different gradient functions across the lamellae of the IVDs using sinusoidal elastic modulus functions. All the constructs were designed to have effective elastic moduli of around 350 MPa, which we calculated using Equation 1 under the assumption that IVDs behave like systems of parallel springs. Only the last two types of IVDs included an FG transition zone (Figure 4E). After manufacturing these specimens and testing them under quasi-static compression, we compared their actual elastic moduli, which were *E*_*D*1_ = 384.3 MPa, *E*_*D*2_ = 332.6 MPa, and *E*_*D*3_ = 322.1 MPa for the first to the final design, respectively. Since the elastic properties estimated with the FEM simulations were highly similar for all the designs (*i.e., E*_*D*1,*FEM*_ = 367.5 MPa, *E*_*D*2,*FEM*_ = 357.1 MPa, and *E*_*D*3,*FEM*_ = 359.3 MPa), we attributed the variations in the measurements to the overestimations that the corrected model yields for lower *ρ* values. Integrating a model correction based on the residual values of the co-continuous model may improve the precision of the designs and is suggested to be performed in future studies. Implementing a machine learning modeling approach may further minimize these errors. However, such a methodology would generally require a large number of experiments and simulations [49] and could betray the purpose of offering a simple and practical model.

The toughness values measured for both graded designs (*i.e*., D2 and D3) were ≈ 2.4 times higher than that of the non-graded design (*U*_*D*1_ = 7.79 MPa, *U*_*D*2_ = 18.86 MPa, and *U*_*D*3_ = 19.05 MPa). The lower toughness of the non-graded design was caused by the sudden separation of the AF from the NP due to their stiffness mismatch (Figure S3D of the supplementary document), leading to a critical stress drop at ≈ 27% strain. In contrast, the graded designs cracked around the annulus fibrosus but did not show critical separation between their phases, which resulted in their continuous hardening. These outcomes support the suitability of the approach chosen for implementing FG in the design of IVDs to improve their toughness. Moreover, these experiments further corroborated the property-by-design approach proposed in the current study that allows for the free adjustment and improvement of the mechanical properties of biomimetic structures for different applications.

### 2.4. Regulating cell behavior

We then moved to a smaller length scale and assessed the possibility of using soft-hard transition functions to regulate the behavior of living cells. More specifically, we hypothesized that we could use the local variations in the elastic modulus of the substrates created through the rational arrangement of micro-bricks to regulate the morphology and function of cells. The use of UV-curable photopolymers in combination with voxel-based additive manufacturing techniques in the biomedical field has been so far limited because the biocompatibility of commercially available UV-curable photopolymers has only been assessed for a few cell types [50,51]. One factor limiting the extensive use of these materials is the adverse effects their leachates have on cells [52,53]. Therefore, prior to the direct seeding of cells on these materials, we assessed the cytotoxicity of materials by exposing hBMSCs and cells from a murine preosteoblast cell line (MC3T3-E1) to the material leachates (Section S4ii of the supplementary document). We did not observe a substantial number of dead cells in any of the conditions considered here. The live/dead images, however, showed limited proliferation of both MC3T3-E1 and BMSC (Figure S4B-C of the supplementary document). The leachates from the soft material (*i.e*., MED625FLX) inhibited proliferation more than those from the hard material (*i.e*., VeroClear) (Figure S4D-E of the supplementary document). For the direct cell cultures, a series of surface treatments consisting of grinding, protein coatings, and the combination thereof were first tested to improve the adhesion of the cells to the 3D printed soft-hard substrates (Sections S4iii, S5, and Figure S5 of the supplementary document). We found that a combination of grinding (SiC abrasive paper, grain size = 5 μm) followed by fetal bovine serum (FBS) protein coating was the most efficient way to improve the cell adherence to the substrates for both hBMSC and MC3T3-E1 cells (Figure S5D-E of the supplementary document).

We first investigated the effects of the hard, soft, graded, and non-graded specimens on the morphology of hBMSC (*i.e*., the area covered by each cell) after one day of direct culture (Figure 5A-B). The cell area on the non-graded soft-hard specimens and at the extremes of the graded specimens was similar to that of their respective monolithic materials, with a clear interface observed between the hard and soft phases on the non-graded specimens (Figure 5B). The mean cell area at the center of the FG specimens was an intermediate value between the values observed for the hard and soft phases, while the values corresponding to the soft and hard extremes were similar to those observed for the non-graded specimens. This finding should be interpreted taking into account the fact that the chemical leachate compositions of the graded and non-graded specimens were the same, meaning that the factor affecting the cell behavior is likely to be local.

**Figure 5.**
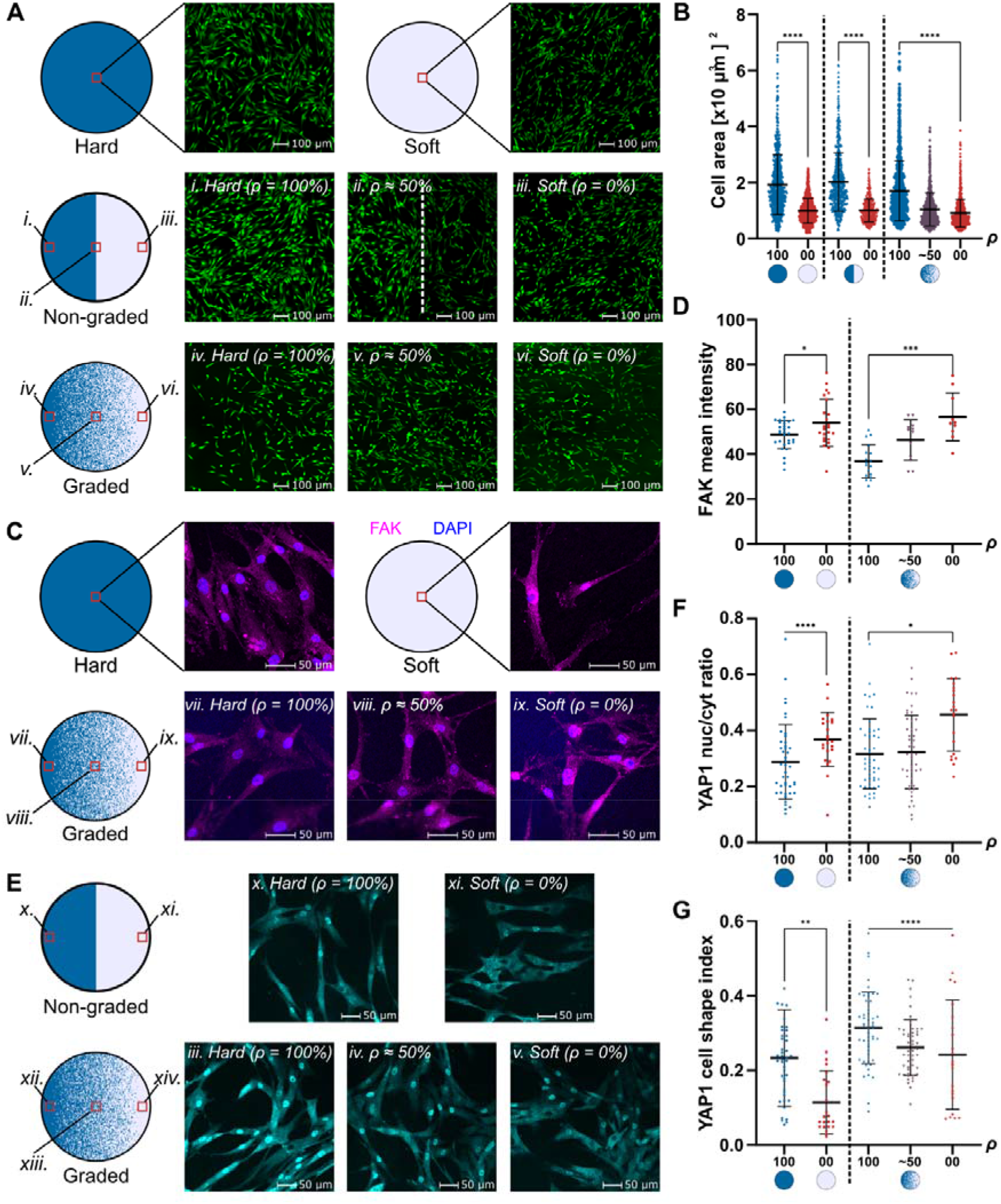
A) The live/dead images corresponding to the monolithically hard and soft, non-graded, and graded specimens. B) A scatter plot indicating the surface area of individual cells at the specified locations across the specimens. An intermediate average surface area value was present at the center of the graded specimens. C) The FAK immunofluorescence staining of the cells seeded on the monolithic and graded specimens. D) A scatter plot depicting the mean FAK intensity signal. An intermediate level (greater than the one for the hard material and less that of the soft material) was present at the center of the graded specimens. E) Representative images of YAP1 (cyan, visualized via immunofluorescent staining) of the cells adhering to the non-graded and graded specimens. F) and G) Quantification of the YAP1 nuclear/cytoplasmatic ratio and cell shape index of the cells adhering to the specimens. The data presented in each scatter plot indicates the value per single cell. Unpaired *t*-tests with the Welch’s correction were performed to compare the ranks of the extremely hard and soft results. The significance of each comparison is marked with *, **, ***, or ****, which correspond to *p* <0.05, 0.01, 0.001, and 0.0001, respectively.

To understand whether the adhesion to the different substrates would result in differential mechano-response, we evaluated the expression of focal adhesion kinase (FAK) by the hBMSC seeded on the graded and monolithic specimens through immunofluorescence staining (Figure 5C). FAK is part of integrin-mediated signal transduction and participates in the formation of focal adhesions between the cells and the substrate [54–57]. The FAK signal was more homogeneously distributed within the cells seeded on the hard material, implying a more uniform formation of focal adhesions within the hard substrate than in the soft one. As for the soft phase, the FAK signal was more intense and was concentrated around the cell nuclei (Figure S6 of the supplementary document). Once again, the use of a graded substrate led to an intermediate level of FAK expression at the center of the specimens (Figure 5D). The mean intensity of FAK on the graded specimens increased gradually from the hard extreme to the soft one. Even though the regulation of FAK expression at the protein level cannot be easily attributed to single mechanical cues of the substrate, these results provide some insight into the potential effects of such soft-hard interfaces and the role of FG. More decisive conclusions, however, can only be drawn with a more thorough investigation in future studies.

Another fundamental factor in mechanosensing and mechanotransduction pathways is the Yes-associated Protein/transcriptional co-activator (YAP1/TAZ) factor [58]. Cell adhesion to substrates results in the assembly of actin fibers, which then transfer the cytoskeletal tension to the nuclei, opening mechanosensitive channels [59]. This process, in turn, allows YAP translocation to the nucleus with enhanced nuclear translocation of YAP corresponding to increased tensile forces. We, therefore, evaluated the presence of this factor and assessed if changes in the mechanical properties of the specimens regulate the nuclear translocation of the hBMSC that were seeded on different types of specimens. The cells seeded on the non-graded specimens showed a different response to the hard and soft materials (Figure 5E). The cells seeded on the regions made of the hard phase had higher YAP1 nuclear to cytoplasmatic ratios than those seeded on the soft phase (Figure 5F). On the graded specimens, the nuclear to cytoplasmatic ratio increased with the presence of the hard phase, although variations existed between cells. The observation that nuclear to cytoplasmatic signal ratio is higher for stiffer materials than for more compliant ones has been reported in the literature [60], which corroborates our results. Furthermore, the YAP staining signal in the regions with mostly hard material was significantly different from those of mostly soft material for both the non-graded and the graded specimens (*p* < 0.05). Similarly, the cell shape index was, on average, lower for the cells seeded on the soft material than those seeded on the hard material for both graded and non-graded specimens (Figure 5G). This observation indicates that the cells residing on the soft material have a more circular shape than those on the hard material. Previous studies have shown that cells exposed to substrates with different stiffnesses tend to migrate toward regions with higher stiffness [61]. It could, therefore, be the case that the cells adhering to the central region of the graded specimens preferentially attach or locally migrate to the stiffer substrate, resulting in the mechanoresponse not being fully correlated with the local ratios of both phases. Nevertheless, the differences between the hard and soft phases in terms of YAP translocation to the nucleus of the cells suggest differential activation of mechanosensitive pathways, which have been shown to play a key role in controlling cell behavior, including growth, proliferation, and differentiation [54,58,62].

## 3. Conclusions

We developed a modified version of classic co-continuous models originally derived for particle-reinforced composites. These models are aimed at establishing a direct relationship between the arrangement of micro-bricks and the macroscale elastic behavior of multi-material 3D printed specimens with voxel-level FGs between their soft and hard phases. Using these models, FGs can be designed at the micro-brick level given the target function describing the variation of the elastic properties between the hard and soft materials. Our experiments and computational models indicated a high degree of correlation between the model-based designs and the actual elastic properties (*R*^2^ > 90%) of such FGs as characterized by both nanoindentation and quasi-static tension tests. We then applied the developed model to design complex biomimetic systems (*i.e*., knee ligaments and IVDs) with pre-programmed variations of elastic properties between their soft and hard phases. The biomimetic specimens incorporating FGs were at least 130% stronger and 140% tougher than their non-graded counterparts, indicating improved load transfer at their soft-hard interfaces. At the cell scale, our experiments supported the hypothesis that cell behavior can be guided by the selective deposition of hard and soft phases within a transition region. Our results, therefore, pave the way for the application of graded soft-hard interfaces fabricated by voxel-level 3D printing to various areas within biomedicine (*e.g*., regenerative medicine and implantable medical devices). Future studies should focus on the characterization of the anisotropic response of soft-hard micro-brick arrangements by canvassing the space of all possible permutations of soft and hard micro-bricks. Moreover, more extensive studies should be performed to better understand how the arrangement of micro-bricks influences the mechanoresponse of cells. Finally, there is a need for more cytocompatible UV-curable photopolymers than can be used with the existing printers to create arbitrarily complex soft-hard interfaces at the voxel level.

## 4. Materials and methods

### 4.1. 3D printing

We fabricated all the specimens through multi-material poly-jet 3D printing (ObjetJ735 Connex3, Stratasys^®^ Ltd., USA). The resolution of the printer (*i.e*., 600×300 dpi in layers of 27 μm) enables a minimum micro-brick size of 42×84×27 μm^3^. The material deposition was controlled using a stack of binary images, which provided a voxel-by-voxel description of the deposition coordinates of both phases. The white bits within each of these stacks represented the location where the 3D printer created each type of micro-brick. We prepared the binary images with MATLAB (R2018b, Mathworks, USA) and processed the prints with GrabCAD Print (Stratasys^®^ Ltd., USA). For most of the mechanical experiments, the hard and soft phases were made from the UV-curable photopolymers VeroCyan™ (RGD841, Stratasys^®^ Ltd., USA) and Agilus30™ Clear (FLX935, Stratasys^®^ Ltd., USA), respectively. The biocompatible MED625FLX™ (Stratasys^®^Ltd., USA) was used as the soft phase in a single nanoindentation specimen with a linear *ρ* gradient and for all the biological experiments. Further details regarding the fabrication process are presented below.

### 4.2. Nanoindentation

#### Specimen design

To create the FGs, we discretized their *ρ*(*x*) functions across the printing direction (*x*-direction, Figure 1A) at the maximum voxel resolution (*i.e*., 42 μm/voxel). For each of the 355 points of the *ρ*(*x*) function, we calculated the total number of hard micro-bricks (*n_H_*) required to achieve their respective *ρ*(*x*) value (*i.e., n_H_*(*x*) = *ρ*(*x*) × *n_layer_, n_layer_* = 177 × 740 voxels^2^) and randomly distributed them over the micro-bricks with the same *x* coordinate. We projected the resulting design at the center of cubic geometries (25×25×20 mm) with the hard phase bounding the FGs. Additionally, the final 1 mm of every design was assigned with *ρ* = 0 %, which served as a reference for the nanoindentation procedure. The shape of the initial FG was a linear function of *ρ* (*ρ_lin_*(*x*) = –*x*/*L_G_* + 100), which was printed twice: once with Agilus Clear as the soft phase and the other with MED625FLX as the soft phase. These specimens were used for material characterization. Later, we defined three specimens with different elastic modulus functions. Their shapes were linear (*E_lin_*(*x*) = –(*E_H_* – *E_S_*)*x*/*L_G_* + E*_H_*) step-wise (*i.e*., *E_step_*(*x*), similar to *E_lin_*(*x*) but discretized in nine steps), and sigmoidal (*E_sig_*(*x*) = (*E_H_* – *E_S_*)/(1 + exp(*d*(*x* – *L_G_*/2))) + *E_S_*, *d* = 8/9 mm^-1^), all with a gradient length of *L_G_* = 14.8 mm. These latter specimens were used for validation. We used a water jet system (Genie 600, Gemini Cleaning Systems, UK) at 12 bar to remove the support material from the specimens.

#### Nanoindentation experiments

We used a TI 950 Triboindenter (Bruker, US) with a diamond conospherical probe with a tip radius of 20 μm to perform the nanoindentation experiments. We followed a previously-described polishing and nanoindentation protocol [32,43]. The nanoindentations were performed in a grid of 33 points along the *x*-direction and 10 points along the *y*-direction, yielding 330 experimental data points per FG. For each FG, the initial position of the nano-indenter was placed at the edge between the regions with only soft and only hard micro-bricks. The distance between successive test points was 500 μm in both directions. Whenever the pull-off forces were >5% of the maximum load, we obtained the reduced elastic modulus (*E*(*r*) of each point using the JKR model [63]. For cases where the pull-off forces were <5% of the maximum load, we used the Oliver-Pharr model [64]. Finally, we calculated the associated elastic moduli (*i.e*., *E*(*x*) ≈ *E_r_*(1 – *v*(*x*)^2^)) by assuming the Poisson’s ratio along the gradient (*v*(*x*)) to be described by the rule of mixtures between the hard and soft phases (*i.e*., *v*(*x*) = *v_H_ ρ*(*x*) + (1 – *ρ*(*x*))*v_S_*, *v_H_* = 0.4, *v_s_* = 0.495). For the linear *ρ* FG results, we compared the resulting elastic modulus functions against predictions made by several existing models for particle reinforced composites, including those proposed by Nielsen [44], Counto [45], and both simplified co-continuous models proposed by Davies (*i.e*., power and logarithmic models) [46]. The comparison with these equations, which are described in detail in Section S1 of the supplementary document, allowed us to obtain the most accurate model that fits our data. For obtaining the *α* value of the modified co-continuous model (Equation (1)), we performed a bisquare non-linear regression between all the available elastic moduli of the linear *ρ* function and their corresponding values of the hard micro-bricks volume fraction. For the linear, step-wise, and sigmoid *E*(*x*) functions, we calculated the coefficients of determination (ordinary *R*^2^ values) between the measured data and the designed functions to validate the accuracy of the selected characterization model.

#### FEM simulations of the nanoindentation experiments

We used a commercial software suite (Abaqus Standard v.6.14, Dassault Systèmes Simulia, France) to perform the FEM simulations of the nanoindentation experiments. Each model was built using a grid of representative volumetric elements (RVEs) taken from 39 positions along the *x*-direction and 9 positions along the *y*-direction (Figure 1B). After a mesh convergence study (Section S2 and Figure S2A of the supplementary document), each RVE included a matrix of 6×6×6 micro-bricks. Each micro-brick was discretized using a cluster of 6×6×6 linear hexahedral elements (C3D8H). We simplified the indenter probe as a cylindrical rigid body defined as a rigid analytic shell with a radius (*R_p_*) of 6.25 μm. We performed four simulations per RVE to account for the nano-indenter position during the tests. In these simulations, the indenter was randomly positioned above the centermost half of the mesh. This process resulted in 1404 simulations per FG. We defined the elastic properties of every hard (*i.e*., *E_H_* = 2000 MPa, *v_H_* = 0.4) and soft (*i.e*., *E_S_* = 0.87 MPa, *v_s_* = 0.495) element based on the NI experimental results. The Poisson’s ratio of both materials and the elastic modulus of the soft phase were obtained from the quasi-static tensile tests performed on monolithic materials (Figure S2B-C of the supplementary document). We constrained all the degrees of freedom of the bottom and lateral regions of every mesh. For the probe, we prescribed an indentation depth (*h*) of 1 μm and recorded its respective reaction forces (*RF*) for each simulation. We calculated the reduced elastic modulus 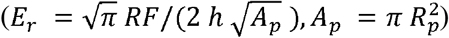 and used the Poisson’s ratios of the RVEs (*i.e*., *v* = *v_H_ρ*(*x*) + *v_s_*(1 – *ρ*(*x*))) to estimate the elastic modulus (*E* = *E_r_* (1 – *v*(*x*)^2^)) for every simulation.

### 4.2. Tensile tests

#### Tensile mechanical tests

First, we prepared monolithic specimens made of only hard and only soft micro-bricks according to the description of type IV specimens in ASTM D638-14 standard [65]. These specimens allowed us to characterize the elastic properties of these materials when loaded in the printing direction. Furthermore, we designed four elastic moduli FGs to validate the accuracy of the corrected characterization models under quasi-static tensile conditions. The shapes of the FGs were a power-law (*i.e*., *E_pow_*(*x*) = (*E_max_* – *E_min_*)((*L_G_* – *x*)/*L_G_* } + *E_min_*, (*i.e*., *E_lin_*(*x*) = –(*E_max_* – *E_min_*)/*L_G_* + *E_max_*) step (*i.e*., *E_step_*(*x*), similar to *E_lin_*(*x*) but discretized using 9 equally spaced steps), and sigmoid (*E_sig_*(*x*) = (*E_max_* – *E_min_*)/(1 + exp(*d*(*x* – *L_G_*/2))) + *E_min_,d* = 8/9 mm^-1^), all with *L_G_* = 12.2 mm. We symmetrically projected these FGs onto the gauge region of the specimens, with their centermost 8.13 mm defined by *E_min_* (Figure 3A). We defined *E_max_* = 1750 MPa and *E_min_* = 75 MPa for all these FGs. The *α*(*x*) functions were obtained using Equation (2) with *α* = 2, *E_S_* = 0.87 MPa, and *E_H_* = 2650 MPa (*E_H_* and *E_S_* were obtained from tensile tests performed on monolithic specimens). We manufactured each design threefold and removed the support material using a water jet system (Genie 600, Gemini Cleaning Systems, UK) at 12 bar. We tested the specimens using an LR5K mechanical testing machine (LLOYD, USA) with a 5 kN load cell at a rate of 2 mm×min^-1^. We recorded the local deformations of the specimens during the tests using a DIC system (Q-400 2× 12 MPixel, LIMESS GmbH, Krefeld, Germany) that captured the surface of the specimens with a frequency of 1 Hz. These measurements required applying a black dot speckle pattern over a white paint background on each specimen. We calculated the first principal (true) strain distributions with the Instra 4D v4.6 (Danted Dynamics A/S, Skovunde, Denmark) software and the DIC measurements. Furthermore, we generated the general stress-strain curves across time (*t*) using the engineering stress (*σ*(*t*) = /(*t*)/*A*_0_, *A*_0_ = 32.512 mm^2^) and engineering strain vectors, *ϵ*(*t*), measured using a digital extensometer within Instra 4D. From these curves, we measured the general elastic modulus of each tensile test specimen, *E_G_*, from the slope of a fitted polynomial of order one. To calculate the elastic modulus along the *x*-direction, *E*(*x*), we fitted polynomials using the average longitudinal (engineering) strains (*ϵ_avg_*(*x*, 0) of each *x*-position and the engineering stress vectors. All the slopes were obtained between stresses of 0.2 and 15 MPa. We then averaged the *E*(*x*) results for three repetitions of each FG and calculated the corresponding coefficients of determinations (ordinary *R*^2^ values) between the experimental results and the FG designs. We further validated this method of measuring *E*(*x*) by calculating the effective elastic modulus 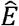 of each test using the equivalent equation for the systems of linear springs 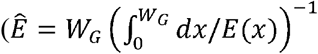, where *W_G_* is the total length of the DIC recording region) and by comparing them to the elastic moduli measured from the general stress-strain curves (*E_G_*).

#### FEM simulations of the tensile tests on FG specimens

We modeled one-half of the gauge section of the tensile test specimens with a cross-section of 24×24 micro-bricks and discretized them with linear hexahedral elements (C3D8H). The resulting meshes consisted of 235008 elements. Given these considerations, we prescribed symmetric boundary conditions at the symmetric end (*i.e*., at the center plane of the soft region) of the design and at two of its lateral surfaces. We also prescribed a uniaxial displacement of 0.26 mm at the hard end of the mesh (equivalent to 1.5% uniaxial strain) while restraining its remaining degrees of freedom. The elastic properties were defined using the tensile test data obtained using the monolithic specimens (*i.e., E_H_* = 2650 MPa, v = 0.4 for the hard phase and *E_S_* = 0.87 MPa, *v_S_* = 0.495 for the soft phase). After processing the simulations, we extracted the reaction forces at the hard end of the FG and the main principal strains at the centroid of every element of the mesh. We performed the same procedure described for the experimental tensile tests to obtain the estimated *E*(*x*) functions of every FG and to calculate their coefficients of determination (ordinary *R*^2^-values) *vs*. the designed functions.

### 4.3. Design, testing, and FEM simulation of knee ligament systems

The geometry of the knee-ligament system was adapted from an open-source CAD database [66]. It consisted of a femur, a tibia, and a fibula (153 mm long) with the respective anterior cruciate (ACL, 15 mm long), posterior cruciate (PCL, 20 mm long), lateral collateral (LCL, 30 mm long), and medial collateral (MCL, 35 mm long) ligaments (Figure 5A). From this assembly, we generated two designs. The first one had FGs between the bone-ligament interfaces. The second design, which worked as our control, was not graded and had abrupt material transitions. The material assignment of both structures was generated with MATLAB (R218b, Mathworks, USA). The FGs of the first design had a sigmoid transition function with *L_G_* = 1.5 mm. To attach the structures to the tensile testing machine, we integrated three cylinders between the femur and tibia regions, which we cut before testing. After removing the support material with a water jet (Genie 600, Gemini Cleaning Systems, UK) at 12 bar, we applied a black dot speckle pattern to a white paint background to measure the system’s deformations with the DIC system. We tested the assemblies under the same conditions as the tensile tests. Furthermore, we built FEM models of both test configurations after reducing their voxel resolution to greyscale RVEs covering 6×6×6 of the original voxels. We discretized each RVE as a single element (C3D8) and assigned its mechanical properties based on its average *ρ* after using Equation (1). We constrained all the degrees of freedom at the bottom surface of the tibial bone. Similarly, we constrained all the degrees of freedom at the top surface of the femur mesh except for the vertical displacement, which was defined as 1 mm. We compared the resulting strain fields of these simulations with the results of the DIC measurements.

### 4.4. Design, testing, and FEM simulation of graded IVDs

We based the dimensions of the IVDs on the L4L5 disc [67] and generated the design with SolidWorks 2021 SP2.0 (Dassault Systèmes, France). The major axis was 49.7 mm, with a minor axis of 31.83 mm and a height of 8.42 mm. After voxelizing the design using MATLAB R2018, we generated three different IVD systems. These IVDs had different sinusoidal FG functions that connected the AF and NP regions (Section S3 of the supplementary document). To discretize these functions, we partitioned the IVDs into concentric lamellae and assigned their respective *ρ* value. The three designs were calculated to have an equivalent elastic modulus of 350 MPa after considering the IVDs as systems of parallel springs (Figure 4E). After printing these designs and removing the support material with a water jet (Genie 600, Gemini Cleaning Systems, UK) at 12 bar, we tested them under uniaxial compression with a mechanical testing machine (Load cell = 100 kN, Zwick Z100, Germany) and using a stroke rate of 2 mm×min^−1^. The stresses and strains were calculated from the loadcell readings and the corresponding crosshead displacements. To obtain more accurate elastic modulus (*E_IVD_*) values, we printed additional specimens of each design and tested them with the same LR5K testing machine (5 kN load cell). The local deformations were measured using the above-described DIC system. Based on these results, we defined a digital extensometer around the centermost part of the IVDs to calculate the average vertical strains of each specimen. Then, we obtained *E_IVD_* from the slope of the linear region of the stress-strain recordings (*i.e*., between the stress values *σ* = 3 MPa and 4 MPa). We compared these measurements with the values predicted by the FE models of these designs, which were built using the same discretization conditions as for the knee ligament constructs. However, each RVE had 3×3×3 voxels, and the simulations were performed under a uniaxial compression equivalent to 5% strain.

### 4.5. Cell culture experiments

#### 3D printed specimens

We designed and 3D printed four types of disk-shaped specimens (*i.e*., with a diameter of 9.75 mm and an out-of-plane thickness of 2 mm) to perform the cell culture experiments. Two of these were made of purely hard (*i.e*., VeroClear™, Stratasys^®^ Ltd., USA) and purely soft (*i.e*., MED625FLX™, Stratasys^®^ Ltd., USA) materials. The other two were designed as non-graded and graded multi-material configurations. The non-graded design had a sharp interface at the center of the disk, with one side of the specimen made only of the hard micro-bricks and the other printed from the soft micro-bricks. The graded design had a linear *ρ* FG analogous to the FG of the initial nanoindentation characterizations.

#### Specimen preparation for direct cell seeding

After 3D printing the specimens, the support material was removed from the printed specimens using a water jet (Genie 600, Gemini Cleaning Systems, UK) at 12 bar. Further removal of support material residuals was done by submerging the specimens in isopropanol under sonication (5510, Branson, UK) for 30 min. Thereafter, the specimens were surface treated by grinding and subsequent FBS coatings, as described in Section S5 of the supplementary document.

#### Cell viability and FAK analyses

Human BMSC (Lonza, 19TL155677) were thawed and plated at 6000 cells/cm^2^ in an expansion culture medium containing a basal alpha minimum essential medium (α-MEM,22571), 10% (v/v) fetal bovine serum (FBS, Hyclone), 100 U/mL penicillin, 100 μg/mL streptomycin, and 10 mM 4-(2-hydroxyethyl)-1-piperazineethanesulfonic acid (HEPES), supplemented with 1 ng/mL of fibroblast growth factor 2 (all from Thermo Fisher Scientific). The culture medium was renewed every two days. Upon reaching 80% confluency, the cells were detached from plastic using 0.05% trypsin-EDTA solution (Thermo Fisher Scientific, USA). At passage three, 104 BMSC suspended in 300 μL (33000 cells/mL) were seeded directly onto the specimens and were kept at 37 °C in a 5% CO_2_ incubator for 2 h. Then, the medium was changed to remove the unadhered cells. The specimens were kept in culture for 48 h, with medium renewal after 24 h, and were then harvested for cell viability and FAK analysis.

Cell viability was analyzed using live/dead assays (LIVE/DEAD^®^ Viability/Cytotoxicity Kit, Thermo Fisher, USA). After removing the culture medium and washing the cells two times with Phosphate-Buffered Saline (PBS, ThermoFisher), we stained the live and dead cells using 2 mM ethidium homodimer-1 and 5 mM calcein-AM for 15 min at room temperature. Then, the solution was removed, and the cells were washed twice with PBS. Finally, the cells were imaged with a ZOE fluorescent cell imager (Bio-Rad, The Netherlands).

For the FAK immunofluorescence staining, BMSC were fixed for 10 min with 2% paraformaldehyde and washed twice with PBS. The cells were permeabilized with 0.5% Triton X-100 for 5 min, followed by 1-hour blocking of non-specific binding sites with PBS with 5%v/v bovine serum albumin (BSA) (Sigma Aldrich) at room temperature. The cells were then incubated with FAK primary mouse monoclonal antibody (1:200, AHO1272, Thermo Fisher Scientific) dissolved in PBS with 1% BSA for 1 hour at room temperature. The specimens were subsequently washed three times with PBS and were incubated with a secondary fluorescent goat anti-mouse AlexaFluor647-conjugate (A21235, ThermoFisher Scientific) at a dilution of 1:1000 in the blocking solution of PBS containing 1% BSA together with 300 nM DAPI nuclei staining. After 1 hour of incubation at room temperature, the specimens were washed with PBS and were stored at 4 °C until the images were taken using a confocal microscope (Leica-SP8, Leica, Germany) using a 20× air-dry objective.

#### YAP analysis

Human bone marrow-derived mesenchymal stromal cells (BMSC) isolated from the surplus bone chips from the iliac crest of a donor (age = 9 years, male) undergoing alveolar bone graft surgery were obtained with the approval of the Medical Ethics Committee of Erasmus MC (MEC-2014-16). The cells were isolated through plastic adherence and were expanded in α-MEM, supplemented with 10% v/v FBS, 1.5 μg/ml fungizone, 50 μg/ml Gentamicin (all Thermo Fisher Scientific, Welthon, MA, USA), 25 μg/ml L-ascorbic acid 2-phosphate (Sigma Aldrich, St. Luis, MO, USA), and 1 ng/ml fibroblast growth factor 2 (Instruchemie, Delfzjil, Netherlands) in a humidified atmosphere at 37 °C with 5% of CO_2_ up to passage 4. 104 BMSC suspended in 300 μL (33000 cells/mL) were seeded directly onto the specimens and were kept at 37 °C in a 5% CO_2_ incubator. The samples were kept in culture for 48 h in a culture medium containing α-MEM supplemented with 10% v/v of FBS 1.5 μg/ml fungizone, 50 μg/ml Gentamicin, and 25 μg/ml L-ascorbic acid 2-phosphate.

For the YAP immunofluorescence staining, the cells were fixated using 4% paraformaldehyde (Boomlap, Meppel, Netherlands) for 10 min, were washed twice with PBS, and were kept in PBS at 4 °C until further processing. To stain the cells, they were permeabilized with 0.5% Triton X-100 for 5 min, followed by blocking the non-specific binding sites with PBS supplemented with 1%v/v bovine serum albumin (BSA) (Sigma Aldrich, St. Luis, MO, USA). The cells were then incubated with primary antibody rabbit anti-YAP1 (1: 500, AB52771, Abcam, Cambridge, UK) dissolved in PBS with 1% BSA for 1 hour. A rabbit Igg isotype (X0903, Agilent Technologies, Santa Clara, CA, USA) was used as the negative control. The specimens were washed three times with PBS and were incubated with secondary fluorescent antibody goat anti-rabbit (1:1000, AB150077, ABCAM, Cambridge, UK) dissolved in PBS with 1% BSA for 1 h at room temperature. The nuclei staining was done using Hoechst dye 33542 (1:2000, Thermo Fisher Scientific, Welthon, MA, USA) for 5 min, followed by two washing steps with PBS 1% with BSA. Finally, the specimens were kept in PBS and were stored in the dark at 4 °C until imaging. Images were taken with a confocal microscope (Leica-SP8, Leica, Germany) using a 20× air-dry objective.

#### Image analysis

All the images were processed using Fiji (version 1.53q, a distribution of ImageJ2, USA). The quantification of the mean FAK and YAP signal intensity, as well as the cell surface area and shape index, were performed using an in-house macro for the Fiji software. To analyze the surface area of the cells based on the live/dead images, the borders of the cells were automatically selected based on thresholding. The same process was applied manually for the FAK and YAP signal intensity analyses because the background signal was too high. The cell shape index (CSI) was calculated from the cytoplasm results of every manually selected cell from the YAP analysis. This index was defined as 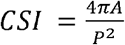, where *A* and *P* represent the area and perimeter of the cell, respectively. A *CSI* value of 1 indicates that the cell is entirely circular, while zero indicates a straight line. The scatter plots of the cell area were created based on 400-500 selected cells for each experimental condition. For the FAK and YAP plots, at least 10 cells were selected to perform the analysis, mainly among the isolated cells that were not overlapping or connected to other cells.

#### Statistical analyses of images

Statistical analyses were performed for the cell surface area, FAK mean signal intensity, and YAP1 nuclei to cytoplasmic ratio using Prism (version 9.4.1, GraphPad Software, USA). Scatter plots were obtained from each corresponding evaluation, where the respective mean and standard deviations were included. For every analysis, we performed unpaired *t*-tests without assuming equal standard deviations (*i.e*., Welch’s correction) to compare the ranks of the results corresponding to the hard and soft materials. We indicated the significance of each comparison with *, **, ***, or ****, which correspond to *p* <0.05, 0.01, 0.001, and 0.0001.

## Acknowledgments

M.J.M acknowledges funding from NWO-Domain Science XS (with code number OCENW.XS22.2.044) and Idea Generator (NWA-IDG) research program with code numbers NWA. 1228.192.206 and NWA. 1228.192.228. This project was partially funded by Open Mind Convergence and the Dutch Medical Delta project: Reg4Med. This research was conducted on a Stratasys^®^ Objet350 Connex3™ printer through the Voxel Print Research Program. This program is an exclusive partnership with Stratasys Education that enhances the value of 3D printing as a powerful platform for experimentation, discovery, and innovation; for more information, contact. We kindly acknowledge Quentin Grossman from the University of Liege for helping with sample polishing prior to nanoindentation. The authors express thanks to Janneke Witte-Bouma and Dr. Eric Farrell from the Department of Oral and Maxillofacial Surgery, Erasmus MC, University Medical Center Rotterdam, the Netherlands, for contributing to the isolation and expansion of cells.

## Authors’ contributions

M.J.M. and M.C.S. designed the research. M.C.S. and R.P.E.V. performed numerical simulations of nanoindentation tests. R.P.E.V., M.C.S., E.L.D., and M.J.M. designed and prepared specimens for nanoindentation tests, uniaxial tensile tests, knee-ligament systems, and intervertebral disc models. A.C., R.P.E.V., and D.R. performed nanoindentation tests and the data analysis thereafter. M.C.S. performed tensile tests of the graded specimens, knee-ligament systems, intervertebral disc, and DIC measurements. M.C.S., M.J.M. S.S., M.K., and M.F. performed specimen preparation for cell culture experiments. S.S., M.K., and M.F. performed direct cell seeding experiments. S.S., M. M., S.L., M.K., and L.E.F performed cell viability and FAK analysis. M.F. and G.J.V.M.O. performed YAP analysis. M.C.S., S.S., M.F., G.T, M.M., S.L., G.J.V.M.O., L.E.F, and M.J.M performed image analyses and data interpretation for cell studies. M.C.S. and S.S. performed statistical analyses. M.C.S., E.L.D., D.R., M.J.M., and A.A.Z performed the data analyses and interpretation for the mechanical test experiments. M.J.M. and A.A.Z. supervised the work. M.C.S., S.S., M.F., M.J.M., and A.A.Z wrote the first draft of the manuscript. All the authors critically revised the manuscript for its intellectual content and approved the manuscript.

