## Supplementary materials for "Rational positioning of 3D printed micro-bricks to realize high-fidelity, multi-functional soft-hard interfaces"

**S1. Characterization of voxel-based particle-reinforced composites**

We evaluated the ability of several models available in the literature to predict the elastic modulus of micro-brick composites. These models included those proposed by Nielsen [1], Counto [2], and two simplified co-continuous models proposed by Davies (*i.e.,* power-based and logarithmic-based) [3] and are formulated as follows:

| Nielsen [1] | $E =\frac{1+A B \rho}{1-\psi B \rho}$  $For: A = k_{e}-1; B = \frac{\frac{E_{H}}{E_{S}}- 1}{\frac{E_{H}}{E_{S}}+A}; k\_e = 2.5$ | (S1) |
| --- | --- | --- |
| Counto [2] | $E = \left( \frac{1-\sqrt{\rho}}{E_{S}} +\frac{1}{\frac{E_{S}\left( 1-\sqrt{\rho} \right)}{\sqrt{\rho}}+E_{H}} \right)^{-1}$ | (S2) |
| Davies (ln) [3] | $E = \exp( \rho\ln\left( E_{H} \right) + (1-\rho)\ln(E_{S}) )$ | (S3) |
| Davies (power) [3] | $E = \left( \rho E_{H}^{1/5} + \left( 1-\rho\right)E_{S}^{1/5} \right)^{5}$ | (S4) |

where $A$, $B$, and $k_{e}$ are model parameters, $\rho$ is the volume ratio of the hard material, and $E_{H}$, $E_{S}$, and *E* are the elastic moduli of the hard, soft, and composite material, respectively. Additionally, we evaluated a modified version of Equation (S3), where we replaced the fixed exponential with a parameter $\alpha$, taking the form:

| Co-continuous (modified) | $E = \left( \rho E_{H}^{1/\alpha} + \left( 1-\rho\right)E_{S}^{1/\alpha} \right)^{\alpha}$ | (S5) |
| --- | --- | --- |

We determined this $\alpha$ parameter through curve fitting using a bisquare non-linear regression algorithm. We evaluated these five models against the NI experimental data (*i.e.,* for AgilusClear and MedFLX625 as soft material) and the estimated FEM nanoindentation data. These evaluations were performed by obtaining the residual plots for the three best-performing models (Figures S1 D-F).

**S2. Mesh convergence test**

We carried out a mesh convergence study prior to the FEM simulations of the NI experiments. More specifically, we evaluated how many micro-bricks per representative volumetric element (RVE) and how many elements per micro-brick were required to accurately model the mechanical behavior of our micro-brick composites. We, therefore, simulated RVEs under the same conditions as described in the main text, where only hard material properties were assumed ($E_{H}$ = 2000 MPa). The selected RVE matrix sizes were 2×2×2, 4×4×4, and 6×6×6 micro-bricks per RVE. Furthermore, we subdivided each micro-brick into arrays of 1×1×1, 2×2×2, 4×4×4, or 6×6×6 C3D8H elements (*i.e.,* equivalent to 1, 8, 64, 216 total elements per micro-brick). Combining these two parameters resulted in 12 discretizations, each of which we simulated 4 times after varying the initial position of the indenter randomly within the top side of the RVE. According to this study (Figure S2A), the error of our simulations was the smallest (*i.e.,* 2.46%) when each RVE consisted of 6×6×6 micro-bricks and each micro-brick was represented by 6×6×6 (*i.e.,* 216) elements.

**S3. Elastic modulus functions for the intervertebral disc designs**

We partitioned the voxelated 3D image of the IVD into 182 concentric lamellae ($n_{l}$ = 182). This partitioning allowed us to define the elastic modulus functions ($E{}_{l}$) in terms of each lamella (*l*) with three distinct regions ($i.e.,$ the annulus fibrosus AF, the FG region, and the nucleus pulposus NP) using the following expression:

$$E_{l}= \left\{ \begin{aligned} \begin{matrix} E_{AF} & ; & l<l_{AF} \\ \frac{E_{AF}-E_{NP}}{2}\left( 1+\cos\left( \frac{l-l_{AF}}{l_{FG}}\pi\right) \right) & ; & l_{AF}\leq l<l_{AF}+l_{FG} \\ E_{NP} & ; & l_{AF}+l_{FG}\leq l \end{matrix} \end{aligned} \right.$$

(S6)

where $E_{AF}$ and $E_{NP}$ are the elastic moduli of the AF and NP, respectively. $l_{AF}$ and $l_{FG}$ are the numbers of the lamellae that define AF and FG regions, respectively. Since the objective was to tune the elastic properties of the IVDs by assuming they behave like systems of parallel springs, we estimated the elastic response of the IVD ($\hat{E}_{IVD}$) as:

$$\hat{E}_{IVD}=\frac{1}{A_{IVD}}\sum_{l=1}^{N_{l}} A_{l}E_{l}$$

(S7)

where $A_{l}$ and $A_{IVD}$ are the surface areas of each lamella and the IVD, respectively. We designed three different IVDs with this approach (*i.e.,* with $l_{G}$ = 1, 32, and 64 lamellae) and tuned the $E_{AF}$ value of Equation (3) to obtain an $E_{IVD}$ of 350 MPa with an $E_{NP}$ of 0.87 MPa and an $l_{AF}$ of 33 lamellae. After this parametrization process, we calculated the equivalent $\rho_{l}$ for each $E_{l}$ using Equation (2) and assuming $E_{S}$ = 0.87 MPa and $E_{H}$ = 2000 MPa.

**S4. Biocompatibility analysis of voxel-based materials**

***i. Materials and Methods:***

*MC3T3-E1 cell preculture*: MC3T3-E1 cells (Sigma Aldrich, Germany) were plated at 4000 cells/cm^2^ in alpha minimum essential medium (α-MEM) supplemented with 10% (v/v) fetal bovine serum (FBS) and 1% (v/v) penicillin-streptomycin. The medium was refreshed every two days. Upon reaching 80% confluency, the cells were detached and used for indirect or direct seeding. To analyze the cytotoxicity of the material leachates, we cultured both BMSC and MC3T3-E1 cells in the extracts of both hard and soft materials. These were prepared by the immersion of the printed, cleaned, and sterilized specimens in 48 well-plates with 1 mL of culture medium. After 24 h of conditioning at 37 °C, the medium from each well was collected, pooled together, and immediately used for cell culture. The medium was changed daily with the medium from the same batch of the conditioned medium that was stored at 4 °C until the test time.

*DNA quantification assay*: 10^4^ cells were initially seeded in 24 well-plates cultured in the hard and soft material extracts (conditioned medium) to assess their effects on the proliferative potential of the cell. After three days of culture, the medium was removed, and the cells were washed and freeze-dried overnight. The cell number was then calculated using the CyQUANT^®^ Cell Proliferation Assay Kit (Invitrogen) according to the provider's protocol. Briefly, the cell lysis buffer was diluted 20X in distilled water and was used to dilute CyQUANT^®^ GR stock solution 400X. 200 µL of the final solution was added to each well, and fluorescence measurements were performed using a microplate reader with an excitation wavelength of 485 nm and emission detection at 530 nm. A reference standard curve was also created separately for both BMSC and MC3T3-E1 according to the providers' protocol for converting the fluorescence values into cell numbers.

*Surface grinding and protein coating:* For the preliminary direct seeding experiments, we tested four different protein coatings: fibronectin bovine plasma (Sigma-Aldrich, USA), collagen (CellAdhere^TM^ Type I Collagen, STEMCELL, USA), medium 10% fetal bovine serum (FBS, Qualified, One shot, Gibco, Thermo Fisher Scientific, USA), and 100% (pure) FBS. The medium composition for each cell type is presented in the main text. Before coating, we placed the specimens in 48 well-plates (Greiner, Bio-One, The Netherlands). For the fibronectin group, we coated each specimen with 500 µL of the fibronectin solution (50 µg/mL in PBS). For the collagen group, we used 500 µL of collagen (STEMCELL, USA) (50 µg/mL in deionized water). For the other two groups, we added 500 µL of culture medium containing 10% FBS or pure FBS to each specimen. After keeping the specimens at room temperature for 2 h, we removed the excess solution from all the specimens.

*SEM imaging*: The cell-seeded specimens were fixated, followed by a dehydration step that consisted of a series of sub-steps: washing with MilliQ water for 10 min, 50% ethanol for 15 min, 70% ethanol for 20 min, and 96% ethanol for 20 min. Then, we soaked the specimens in hexamethyldisilazane (Sigma Aldrich, USA) for 15 min and left them to dry in the open air overnight. A scanning electron microscope (SEM, JEOL JSM-IT100, Japan) was used to acquire the images. The specimens were gold-sputtered prior to SEM analysis.

***ii. Indirect seeding and leachate analysis****:*

We analyzed the cytotoxicity of the leachates obtained from the UV-curable materials. We performed this task by exposing the BMSC and the MC3T3-E1 cells to material-extracted substances for up to 3 days. On days 1 and 3, the metabolic activity of the cells, the number of the cells, and their viability were determined.

The live/dead results showed few dead cells in all the conditions (Figure S4B-C). The culturing of the MC3T3-E1 cells in the extracts from the hard material caused the cells to exhibit 78% of the metabolic activity of the control group on day 3 (Figure S4D). As for the soft material, the viability decreased to 16% on day 3 (Figure S4E). The DNA quantification results indicated that the leachates of the soft material decreased the proliferation potential of the MC3T3-E1 and BMSCs more significantly than the hard material (*p* < 0.01).

***iii. Direct seeding and protocol development:***

The BMSC and MC3T3 cells were seeded directly on the printed substrates (Figure S5). Only a few cells adhered to the surface of the untreated specimens after 24 h (Figure S5B). To enhance the number of adhered cells, we smoothened the surface topography of the printed substrates by grinding them with two different SiC abrasive papers (grain size of 10 and 5 um). In addition, we tested the effects of four different protein coatings (*i.e.,* fibronectin, collagen, culture medium containing 10% FBS, and pure FBS). Among the tested protein coatings, the medium containing 10% FBS increased the number of cells that adhered to the substrate, exceeding the performance of pure FBS. In fact, FBS coating enhanced the cell attachment for both cell types on both materials. For all conditions, fewer cells adhered to the soft material than to the hard one. Applying the grinding process was possible for all the substrate types, even when both phases were non-graded (Figure S5C). After grinding, the cell seeding results showed enhanced seeding efficiency and viability, with the best results obtained after grinding with a fine SiC paper with a grain size of 5 µm. This outcome was further improved when combining grinding with pure FBS coating (Figure S5D-F). Consequently, this surface optimization protocol allowed us to perform additional experiments to evaluate the cell behavior in specimens containing hard and soft materials in graded and non-graded configurations.

**Table S1.** A comparison of the effective elastic moduli found through tensile tests. All values are in MPa. The values calculated using the local response of the DIC images ($\hat{E}$) are compared with the measurements using a digital extensometer ($E_{G}$). The overall ordinary coefficient of determination was *R^2^* = 95.76%

|  | Power-law | | | Linear | | | Step-wise | | | Sigmoidal | | |
| --- | --- | --- | --- | --- | --- | --- | --- | --- | --- | --- | --- | --- |
| $\hat{\boldsymbol{E}}$ | 27.0 | 28.6 | 27.7 | 29.2 | 18.5 | 32.0 | 28.9 | 30.6 | 31.2 | 18.1 | 17.8 | 16.4 |
| $\boldsymbol{E}_{\boldsymbol{G}}$ | 24.7 | 28.1 | 29.3 | 28.2 | 18.8 | 29.1 | 27.2 | 28.8 | 29.8 | 16.0 | 17.4 | 14.9 |

**Supplementary figures**

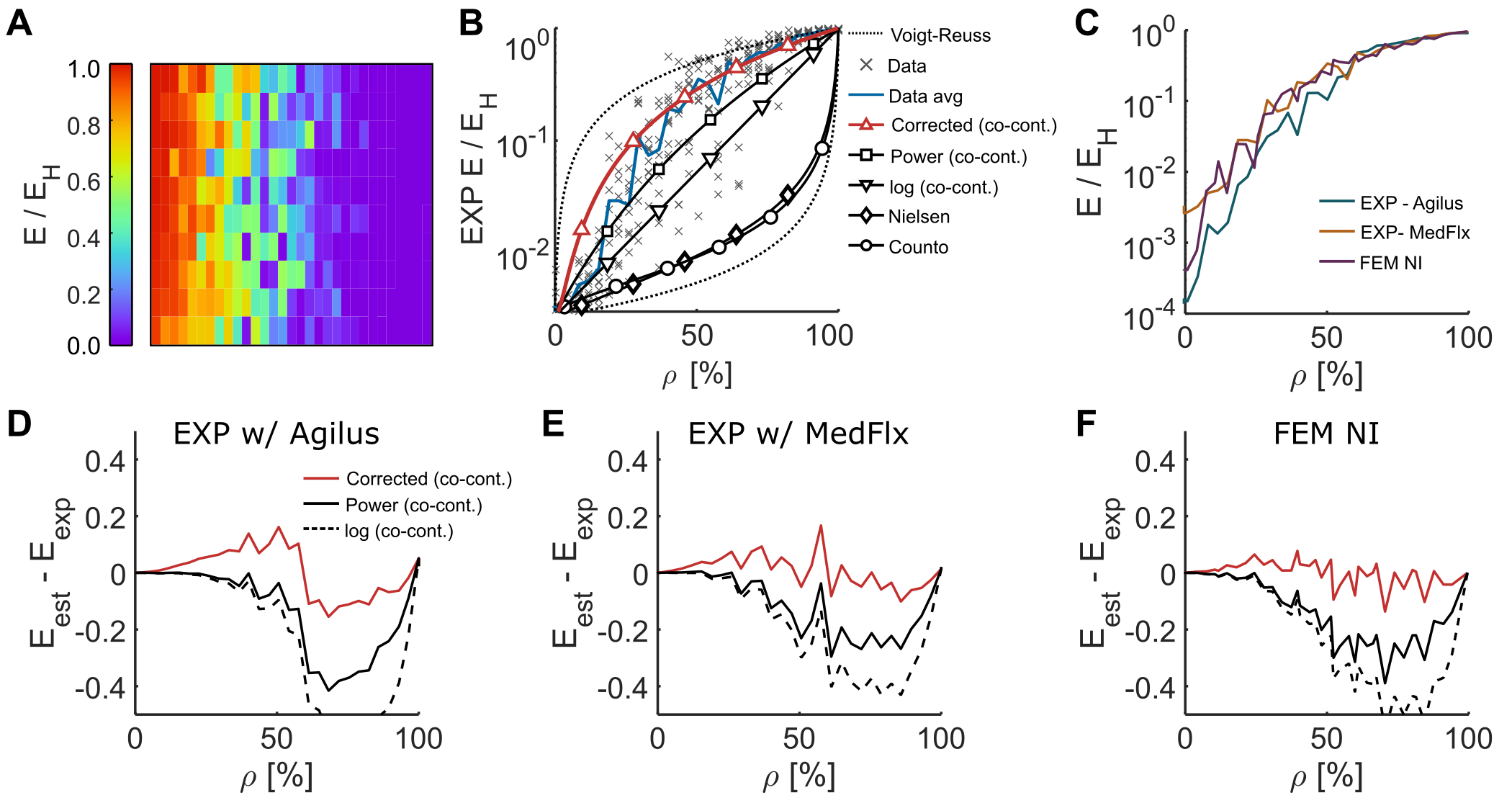

**Figure S1.** A) The distribution and B) average values of the elastic modulus as measured by NI for a soft phase made from MED625FLX^TM^ (Stratasys^®^ Ltd., USA). The models used for comparison are enlisted in the S1 section of this document. C) A comparison between the average response of the experimental datasets and FEM estimations. D-F) The residual plots ($E_{est}$-$E_{exp}$) of the three best-performing models (*i.e.,* Equations S3, S4, and S5) for the NI experiments performed with Agilus30 (D) and MED625FLX (E) as well as for the corresponding FEM estimations (F).

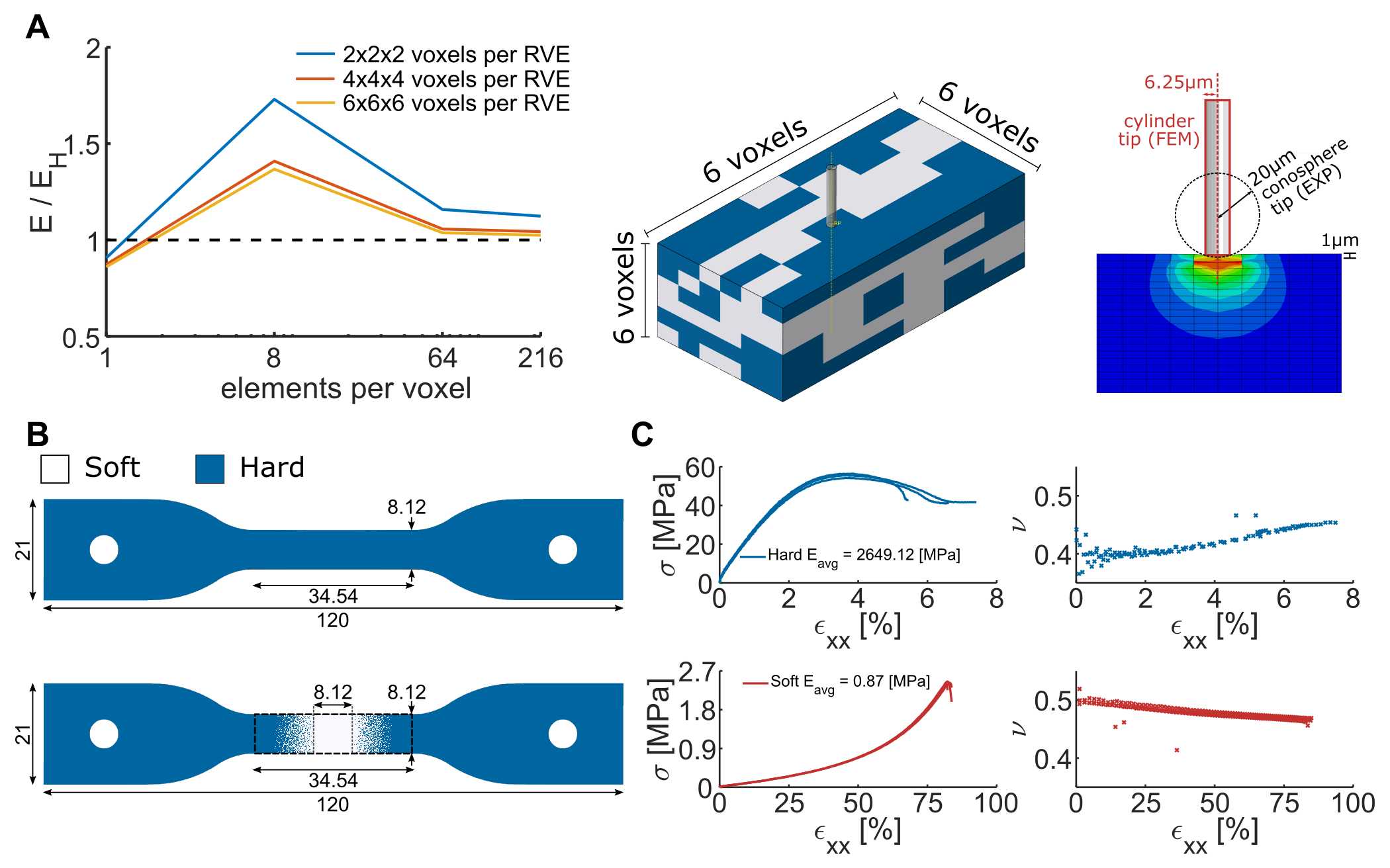

**Figure S2.** A) The results of the mesh convergence study. RVEs consisting of 6×6×6 micro-bricks and 216 linear hexahedral linear elements per micro-cube led to the smallest error when choosing a flat indenter with a radius of 5 µm and an indentation depth of 1 µm. B) the designs of the monolithic specimens used in quasi-static tensile testing (thickness = 4 mm). C) The results of the quasi-static tensile tests performed on the monolithic specimens. We obtained the average elastic modulus of both phases for the specimens printed in the loading direction. The Poisson's ratios of the materials were determined using the DIC system.

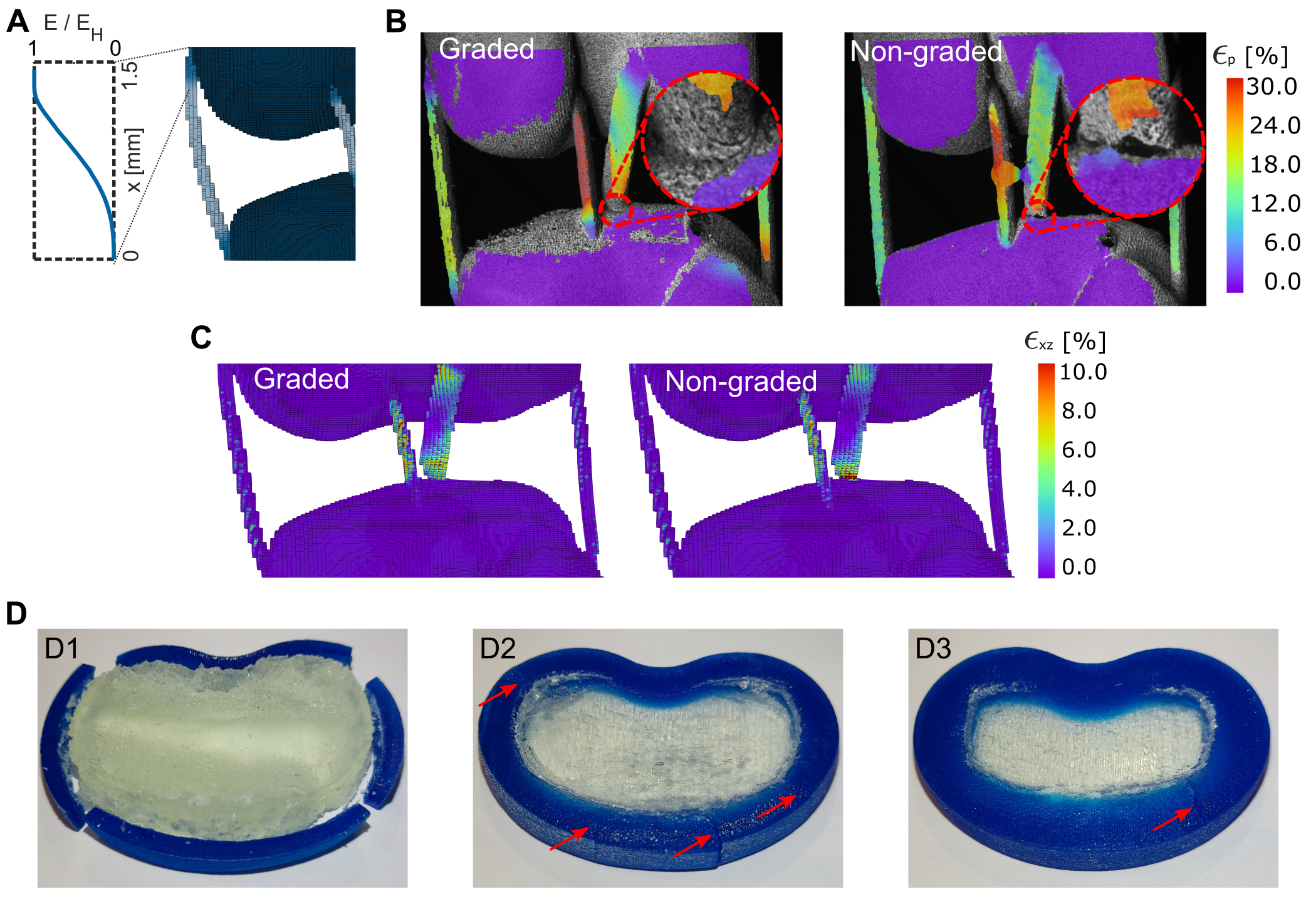

**Figure S3.** A) Representative example for discretizing an FG in the knee-ligament system. A sigmoidal gradient with a length of 1.5 mm was applied to every bone-ligament interface. B) The DIC images of the graded and non-graded knee ligament constructs after 5 mm of deformation. We observed the propagation of a non-critical crack in the non-graded specimen while the graded sample remained fully bonded. C) The distribution of the shear strain ($є_{xz}$) within the graded and non-graded knee-ligament as predicted by the FEM models. These results show the relationship between the strain concentrations present in the non-graded specimens and shear deformations, which are highly reduced when an FG is applied. D) The failure modes of the three IVD designs after the quasi-static compression tests. The gradient-less (D1) design showed a total separation between both material phases, while the graded designs did not fail critically and merely presented non-critical cracks around the annulus fibrosus regions.

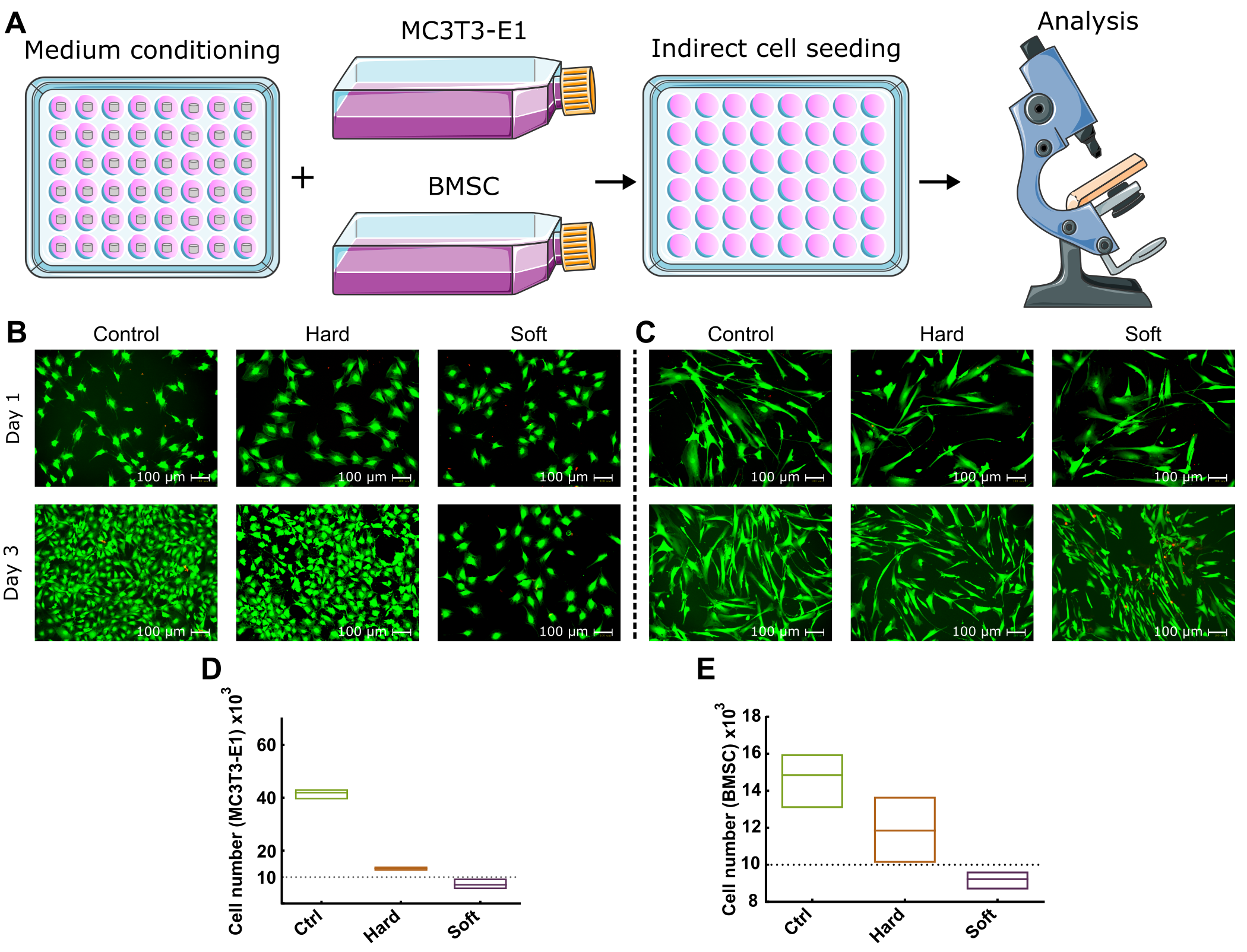

**Figure S4.** The results of indirect cell seeding in conditioned medium. A) A schematic drawing of the workflow used for indirect cell seeding. Each specimen was immersed in a culture medium for 24 h. The collected medium was used to culture the cells in standard well-plates. B-C) The live/dead images of MC3T3-E1 (B) and BMSC (C) after 1 and 3 days of culture in the extracts of the hard (VeroClear) and soft (MED625FLX) materials. Although few dead cells can be detected in all the conditions considered here, the cells cultured in the extracts of the soft material were smaller in number as compared to that of the hard material or that of the control group. D) and E) indicate the number of the MC3T3-E1 cells and BMSC cultured with the extracts.

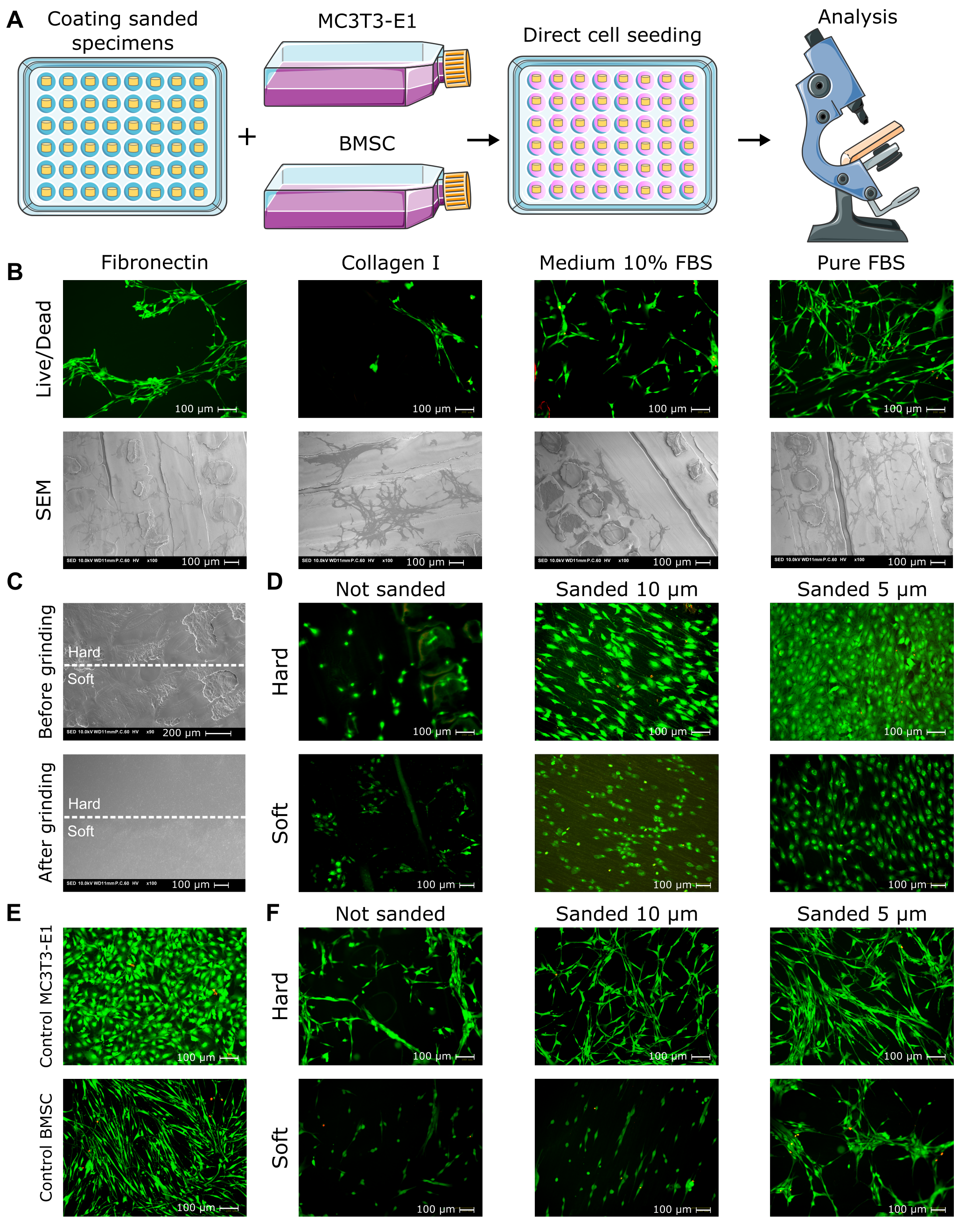

**Figure S5.** Direct cell seeding and the optimization of its protocol. A) A schematic drawing of the workflow for direct cell seeding. B) The live/dead images of the BMSC after 1 day of culture on the specimens with different types of protein coatings. C) The SEM images comparing the surface topography of a non-graded specimen before and after applying the grinding process using a 5 µm SiC paper. D) The direct seeding of the MC3T3-E1 cells cultured on the hard and soft specimens after their surfaces were grounded with SiC papers (5 and 10 µm) and coated with 100% (pure) FBS. The live/dead assay was performed after 24 h. E) The live/dead assays of the control groups (seeded on standard well-plates) for both cell types were performed after 24 h. F) The direct seeding of BMSC on the hard and soft specimens after grinding with 5 and 10 µm SiC papers and coating with 100% (pure) FBS. The live/dead assay was performed after 24 h.

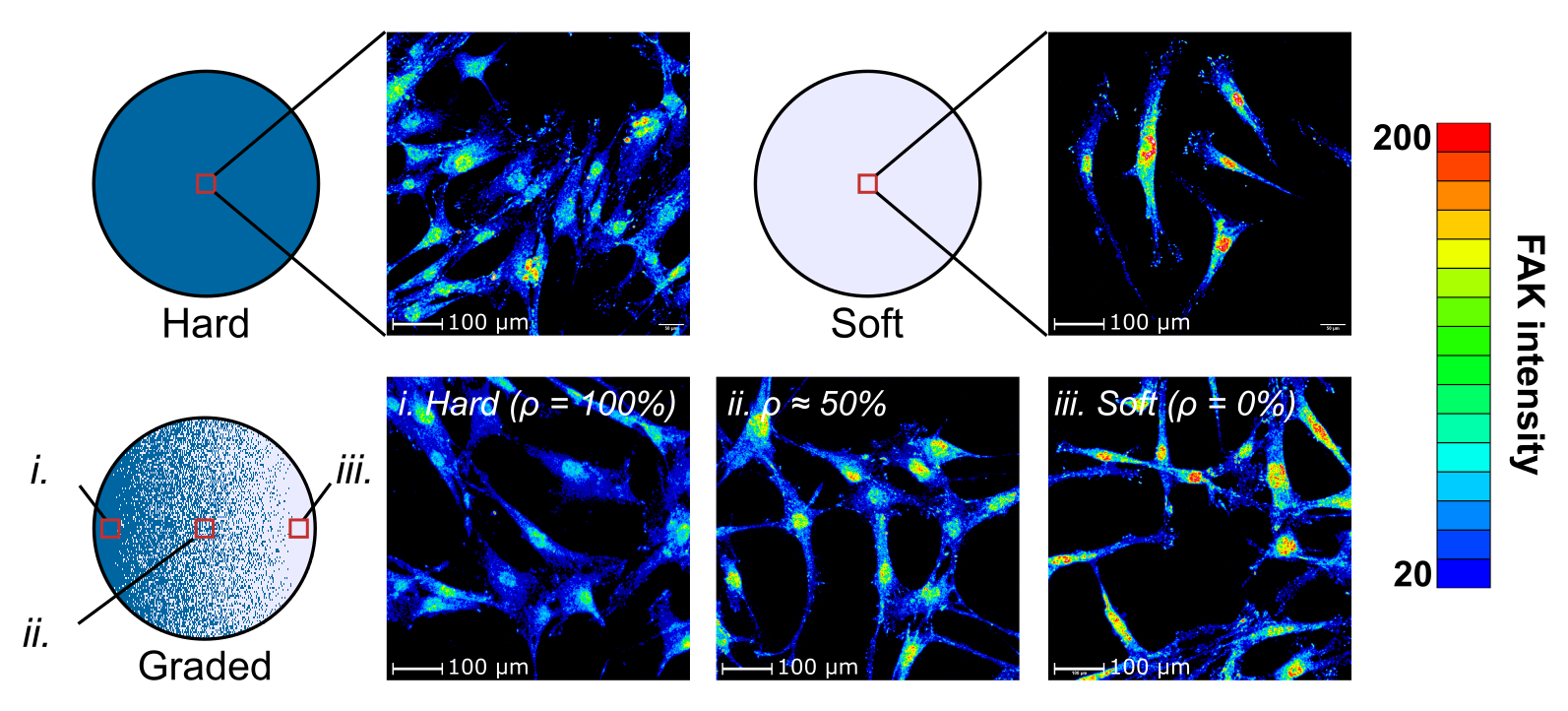

**Figure S6**. The heatmap of the FAK signal. The cells on the hard material (A) exhibited more uniform distributions of the signal, while the cells seeded on the soft material (B) showed a concentrated area around the nuclei where the FAK signal intensity was high. This trend was also observed for the cells cultured on the graded specimens. C) The signal intensity in the regions close to the nuclei increased with the volumetric percentage of the soft material.
